# pH-responsive self-assembled compartments as tuneable model protocellular membrane systems

**DOI:** 10.1101/2022.05.14.491926

**Authors:** Susovan Sarkar, Shikha Dagar, Kushan Lahiri, Sudha Rajamani

**Affiliations:** Department of Biology, Indian Institute of Science Education and Research, Pune 411008, India

## Abstract

Prebiotically plausible single-chain amphiphiles are enticing as model protocellular compartments to study the emergence of cellular life owing to their self-assembling properties. Here, we investigated the self-assembly behaviour of mono-N-dodecyl phosphate (DDP) and mixed systems of DDP with dodecyl 1-dodecanol (DOH) at varying pH conditions. Membranes composed of DDP showed pH-responsive vesicle formation in a wide range of pH with a low critical bilayer concentration (CBC). Further, the addition of DOH to DDP membrane system enhanced vesicle formation and stability in alkaline pH regimes. We also compared the high-temperature behaviour of DDP and DDP-DOH membranes with conventional fatty acid membranes. Both, DDP and DDP:DOH mixed membranes possess packing that is similar to decanoic acid membrane. However, the micropolarity of these systems are similar to phospholipid membranes. Finally, the pH-dependent modulation of different phospholipid membranes doped with DDP was also demonstrated to engineer tuneable membranes with potential translational implications.

## 1. Introduction

Semipermeable boundaries composed of amphiphiles are considered to be crucial for the emergence of cellular life on the early Earth (protocells) ^[1–3]^. Protocellular membranes are assumed to be relatively simpler and composed of single chain amphiphiles (SCAs) ^[2,3]^. Fatty acids (FA) have been studied extensively in this regard because of their prebiotic relevance, ability to assemble into cell-sized vesicles and the capacity to encapsulate solutes ^[2–4]^. Unlike diacyl phospholipids (PL), FA vesicles, due to their unique dynamic nature, possess properties such as high permeability to solutes and increased fluidity, which are crucial to support the emergence and sustenance of protocells ^[3–5]^. FAs can assemble into different structures under aqueous conditions depending upon the surrounding pH. In acidic and alkaline pH conditions, the FAs are predominantly in their protonated and deprotonated states and thus assemble into oil droplets and micellar structures, respectively ^[6]^. FAs assemble into vesicles within a narrow pH range (approximately 7–8.5), which is close to the apparent pKa of their acid head group (negative logarithm of the acid dissociation constant)^[3,6]^. This ability to form vesicles only in a narrow pH regime results in significant shortfalls and limits the environments that would have been suitable for the emergence of protocells ^[4,7,8]^. Previous studies showed that certain prebiotically pertinent reactions such as RNA polymerization ^[9,10]^, nucleotide activation chemistry ^[11]^ and nonenzymatic replication ^[12]^, are all facilitated in acidic pH regimes. Also, the RNA phosphodiester backbone is most stable in the pH range 4-5 ^[10]^. On the other hand, alkaline pH ^[9,10]^ can facilitate amino acid oligomerization ^[13]^ and formose reaction ^[14]^. This emphasizes the FA-based membranes’ disadvantage to support the emergence and sustenance of protocellular life forms.

Towards this, the addition of co-surfactants like fatty alcohols and fatty acid monoglycerides has been shown to stabilize FA-based vesicles in alkaline pH regimes but not in acidic pH ^[8,15]^. Moreover, the archean oceans are proposed to have had acidic pH ^[16]^. This underlines the importance of exploring alternate, prebiotically plausible membrane forming SCAs. Towards this, Powner and Sutherland reported the synthesis of alkylphosphates (aliphatic chains containing phosphate head groups) from fatty alcohols under prebiotic settings ^[16]^. Phosphates is also a major constituent of contemporary plasma membranes and genetic material backbone (DNA and RNA). Traces of short alkylphosphates have also been found in meteorites ^[17]^. Previous studies have reported the ability of different alkylphosphates to assemble into vesicles in narrow acidic pH (around pH 2) ^[18,19]^. Because of the low solubility of alkylphosphates under aqueous and ambient conditions (RT), these assemblies were found to be unstable. Albertsen et al. added co-surfactants with decyl phosphate to overcome this problem and looked at the vesicle forming ability of mixed decyl phosphate (C10) systems ^[20]^. Recently, Gao M et al., investigated the vesicle formation of sodium dodecyl phosphate (C_12_) in neutral pH in *n*-butanol/water mixed solvent to increase its solubility ^[21]^.

Inspired by these studies, we aimed to systematically characterize the self-assembly behaviour of mono-N-dodecyl phosphate (DDP) and discern its membrane’s physicochemical properties. We explored the aggregation behaviour and vesicle formation propensity of DDP from pH 2 to 10 using microscopy, UV-vis spectroscopy and photon correlation spectroscopy. Using steady-state fluorescence spectroscopy, the different properties of DDP membranes, such as micropolarity, order (packing) and fluidity were also determined over varying pH. The effect of the addition of varying ratios of 1-dodecanol (DOH), a prebiotically plausible co-surfactant, on the self-assembly and properties of DDP-DOH membrane systems was also characterized. Our results show that DDP alone can assemble into vesicle over a wide range of pH from 2 to 10. However, the assembly process and the nature of the self-assembled structures are dependent on the surrounding pH. Interestingly, DDP membrane properties were found to be responsive to pH change owing to different protonation states of DDP molecules at a given pH. DDP membrane micropolarity and fluidity increased, with increasing pH, and the membrane packing (order) decreased with increasing pH. DOH was found to modulate the self-assembly behaviour and properties of DDP membranes when doped in varying ratios. DOH decreased the overall membrane packing and stabilized the mixed vesicles under alkaline conditions potentially by providing a H-bonding partner.

Terrestrial hydrothermal hot springs ^[22]^ and deep-sea hydrothermal vents ^[23]^ have been proposed as potential sites for the emergence of early cellular life on Earth. They both are characterized by elevated temperatures (about 50-90°C). However, the high-temperature behaviour of model protocellular membranes remains understudied as predominant studies have been carried out at ambient conditions (room temperatures) ^[1–6,15,20^]. Mansy et. al. reported thermostable membranes composed of decanoic acid-decanol and decanoic acid-decanol-monocaprin (2:1 and 4:1:1 molar ratio, respectively) that were able to retain encapsulated oligonucleotides up to 70°C for more than 1 hour ^[24]^. Galvanized by this, we studied the high-temperature behaviour of both DDP-based and DDP-DOH mixed membranes, at varying temperature (35°C to 65°C) and in different pH conditions. We also compared these membrane systems with other conventional FA-based membranes. The membrane packing was found to decrease with an increase in temperature, suggesting an increase in permeability. The degree of change of this packing parameter was dependent on the membrane composition and the surrounding pH. Apart from this change, our results also indicated morphological changes in the self-assembled structures upon an increase in temperature. Importantly, DDP molecules were also found to assemble into stable bilayers more readily at lower concentrations when compared to FA of the same chain length. On comparing the properties of DDP membranes with different PL and FA-based membranes at pH 8, DDP membranes showed similar micropolarity as PL membranes. Nonetheless, DDP membrane packing was similar to that of FA-based membranes indicating their unique dynamic nature. At pH 4, DDP membranes showed compact packing and low fluidity. The crystalline nature of these membranes at pH 4 was confirmed by the presence of sharp peaks in XRD pattern which were absent at pH 8. Pertinently, this highlighted the tuneable nature of DDP membranes in response to pH and the ability of these compartments to change their property and adapt in response to changes in the immediate environment. Subsequently, we engineered novel tuneable membrane systems by mixing DDP with PL, and also show that this pH-responsive tuneable nature of DDP membranes can also be also extended to the resultant hybrid membranes. Specifically, the addition of DDP to different PL membranes was found to change the membrane order in response to pH. Such mixed PL-DDP membrane systems with tuneable nature can also be useful in synthetic biology applications and for engineering biosensors.

## 2. Self-assembly of DDP and physicochemical properties of DDP self-assembled structures at varying pH

### 2a. pH dependent self-assembly behaviour of DDP

To understand the pH dependent self-assembly behaviour of DDP, the suspension’s turbidity was measured at 600 nm from pH 2 to 10. Aggregates such as vesicles, droplets and crystalline structures would scatter more light than monomers and micellar aggregates because of their large size, leading to an increase in turbidity ^[15,25]^. As shown in Fig 1a and Fig S2, the highest turbidity (1.2±0.08) was observed in pH 2 indicating the formation of large aggregates. The turbidity then decreased moderately in a non-linear manner with increasing pH till pH 8. A sudden decrease in turbidity was observed in the alkaline pH between 9 to 10 (~0.29) indicating a structural transition and potential presence of smaller aggregates such as micelles and monomers. To understand this further we used pyrene, a solvatochromic fluorophore. Upon excitation at 335 nm, the pyrene monomer shows five vibronic peaks in the range of 370-400 nm. The intensity ratio for the first and third peak (I_1_/I_3_) is known to indicate the polarity of the pyrene microenvironment ^[26,27]^. The ratio is highest in water (1.68±0.08) (Fig S3) and it decreases upon partitioning of pyrene into hydrophobic regions. In DDP suspension, the I_1_/I_3_ ratio was observed to be in between 0.89 - 0.85 from pH 2 to 10, denoting the presence of hydrophobic aggregates (Fig S4). The ratio was highest at pH 2 (0.89) and gradually decreased with increasing pH (till pH 8) indicating a decrease in polarity. The ratio again increased significantly at pH 9 and 10, indicating structural transition corroborating with the observed decrease in turbidity.

**Figure 1:**
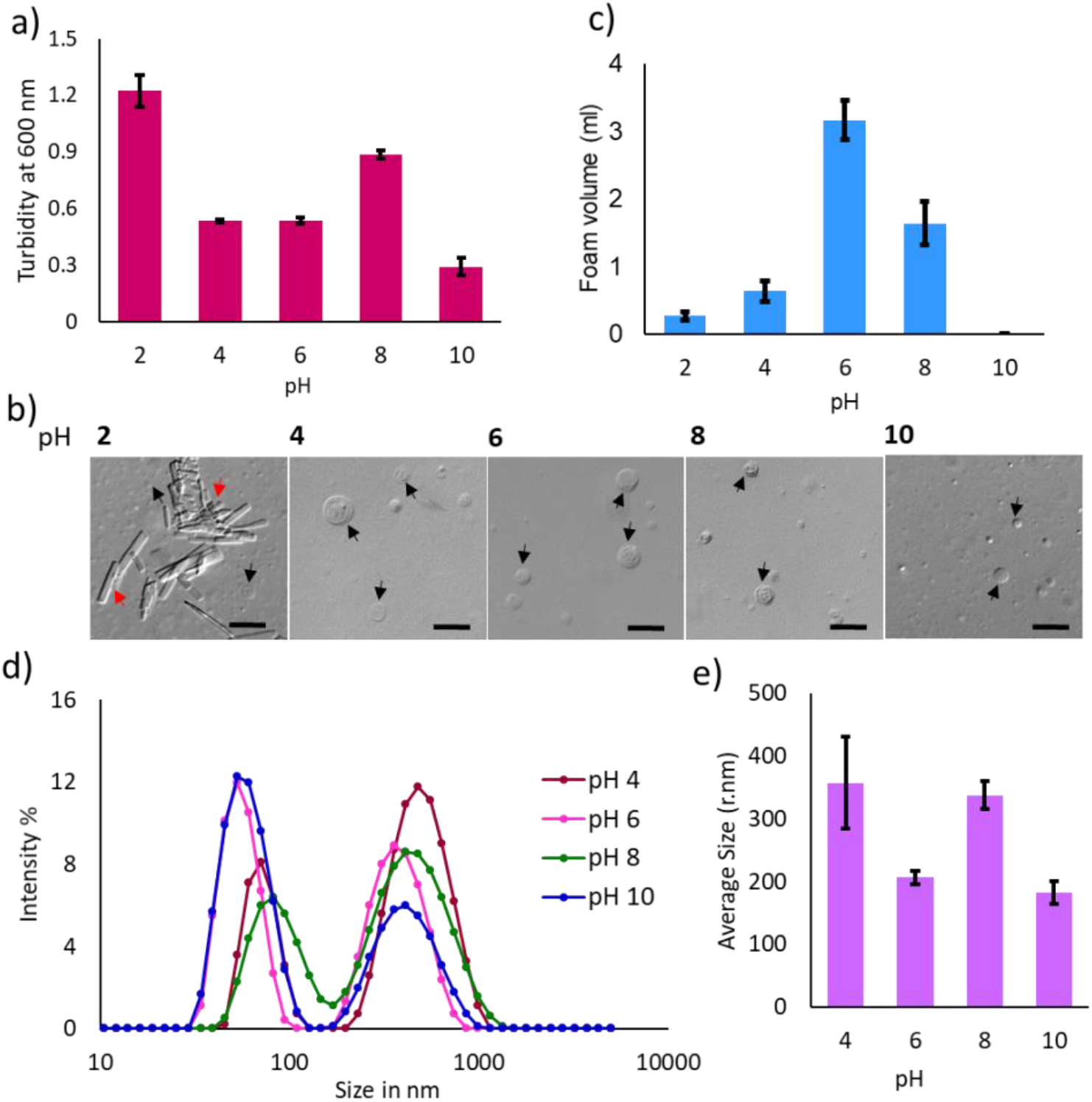
a) The bar plot shows the turbidity measurements of the DDP suspension at different pH (x-axis) at 600 nm and 45°C, N = 3, error bar = SD. b) DIC microscopy images of DDP suspension over different pH. Black and red arrows indicate vesicles and crystalline aggregates, respectively. N = 3, Scale bar = 10 μm. c) Foam volume (ml) measurements (foamability) of DDP suspension at different pH. N = 3, error bar = SD. d) The scatter plot shows the size distribution of the self-assembled structures of DDP suspension at different pH plotted as a function of % intensity of scattered light at 45°C, N = 4. e) Average size (nm) of the population of DDP suspension plotted over different pH, N = 4, error bar = SD.

Next, to investigate the nature of the DDP self-assembled structures at different pH, the suspensions were examined under the microscope. Large vesicles were observed from pH 2 to 10 as shown in Fig 1b and Fig. S5. However, at pH 2, large crystalline aggregates were also observed along with vesicles, explaining the high turbidity. From pH 3 to 10, only large micron-size oligovesicular vesicles were observed. At pH 8, aggregation of vesicular structures was observed that resulted in rosset-like structures (Fig S6). To determine the nature of the self-assembled structures in the suspension, foamability and foam stability assay were carried out at different pH (pH 2, 4, 6, 8 and 10). Foam formation and its stability in a surfactant suspension depend upon gas transfer between bubbles, the flow of liquid in the foam, and film rupture between bubbles ^[28]^. Large assemblies such as vesicles and lamellar structures increase the local viscosity of the suspension hence, decreasing the drainage of liquid from the foam and preventing the transfer of gases between bubbles thereby, stabilizing foam ^[28,29]^. As a result, the foam produced by suspensions containing vesicles is typically larger in volume as compared to micellar suspension ^[29]^. This is because the smaller size of the micelles fails to increase the local viscosity of the suspension and prevent the drainage of the liquid. Further, the repulsion between deprotonated surfactants also prevents the formation of a rigid barrier at the air/water interface. In our experiments, the highest foam volume of 3.1±0.2 ml was observed at pH 6, followed by 1.6±0.3 ml of foam at pH 8, indicating that a higher amount of vesicular structures was potentially present in the suspension at pH 6 (Fig 1c and Fig. S8). No foam was observed at pH 10, confirming the predominance of micelles and monomers in the suspension. Low volumes of foam of 0.2±0.05 ml and 0.6±0.1 ml, were observed in pH 2 and pH 4, respectively. The foam was completely diminished at pH 2, 4 and 8 after 2 hours (Fig S7 and S8). At pH 6, the foam was found to be stable even after 8 hours of incubation suggesting the extent of vesicular self-assembly that was facilitated at this pH (Fig S9).

This behaviour of self-assembly over varying pH can be explained by factoring in the protonation state and the solubility of DDP in water. Previous studies report the p*K*a values of DDP to be 2.85 (p*K*a1) and 7.35 (p*K*a2) as it possesses two proton donating sites ^[30]^. At pH 2, 87.5% of DDP molecules remain in a doubly protonated state [C_12_H_25_OPO(OH)_2_] resulting in low solubility in water. With increasing pH, DDP gets gradually deprotonated hence its solubility increases (Fig. S8). The mono-protonated species C_12_H_25_OP(OH)O_2_^−^ can H-bond by themselves or with the doubly-protonated species (C_12_H_25_OPO_3_^2−^. This can result in the formation of acid-soap complex and thereby facilitating membrane formation as has been observed in the case of FAs too ^[6,31]^. With increasing pH, the amount of mono and di-deprotonated species increases as summarized in table ST1. At pH 4, 93.45% and 6.5% of DDP are in the mono-protonated and doubly protonated state, respectively. 95% of DDP remains in the mono-protonated state with 5% of the di-deprotonated species at pH 6, which results in efficient H-bonding and vesicle formation. At pH 8, the mono-protonated and di-deprotonated species are present in 81% and 19% respectively, reflecting a comparatively more negative charge on the vesicle surface. More than 99% of DDP remains in the di-deprotonated state at pH 10. The presence of two negative charges on the head group would bring about repulsion between the DDP molecules hence, resulting in predominant micelle formation at pH 10.

To investigate the size of the vesicular structures at different pH, dynamic light scattering (DLS) was used. The autocorrelation functions data (g) was found to fit the double exponential model for pH 4, 6, 8 and 10 (Fig S11-S14), because of the varying sizes in the suspension ^[32] [33]^. However, the presence of large precipitating crystalline aggregates observed at pH 2, hindered the DLS measurement of this sample, hence was not considered. Such large precipitating aggregates were observed in the suspension (at pH 2) even after incubation for 5 minutes without any shaking at 45°C (Fig. S15). Upon plotting the scattering intensity as a function of particle size, we observed a bimodal distribution, with a smaller size distribution around 50 to 150 nm and a larger size distribution around 250 to 1000 nm of the self-assembled structures for pH 4, 6, 8 and 10 (Fig 1d). The particles with bigger size contribute more to scattering light compared to smaller size ones. In the case of pH 4 and 8, the scattering intensity contribution from the larger size population was higher as compared to the scattering intensity from that of the smaller size population, with an average size of 356±73 and 337±22 nm, respectively (Fig 1e). At pH 10, the average size of the population was found to be lowest at 183±17 nm, potentially because of the presence of micellar structures.

### 2b. Physicochemical properties of DDP membrane as a function of pH

To better understand the physicochemical properties of DDP membranes, and the effect of surrounding pH on membrane properties, we used different solvatochromic probes. DDP membrane micropolarity at different pHs was measured using nile red (NR), a hydrophobic fluorophore. NR shows an emission maximum at 660 nm in bulk water (Fig S16a). Upon incorporation into a hydrophobic environment, a blue shift is observed in its emission spectrum indicating less water accessibility ^[27,34]^. Studies using PL membranes show that NR localizes in the interfacial region of the membranes ^[35]^. The NR emission intensity ratio of 610 nm to 660 nm (I_610_/I_660_) was measured to gauge the micropolarity of the DDP membrane. Higher the ratio, lesser is the water accessibility on the membrane, thereby, indicating a decrease in micropolarity ^[27,34]^. NR I_610_/I_660_ was found to be lowest (0.2±0.03) in water as shown in Fig S16b. NR showed an emission maximum at 590 nm upon incorporation in the DDP membrane at pH 4 with the I_610_/I_660_ ratio of 2.4±06, indicating the low micropolarity of the membrane (Fig 2a and Fig. S17). From pH 4 to 6, the ratio decreased to 2.2±0.03 with a new emission maximum at 620 nm, indicating increased micropolarity. From pH 6 to 10, micropolarity remained unaltered with an emission maximum at 620 nm (Fig. 2a and Fig. S17). This showed that upon increasing the pH from 4 to 6, the water accessibility on the membrane increases and remains unchanged above pH 6. However, for PL membranes (POPC), the I_610_/I_660_ ratio was found to be constant over a range of pH (4 to 10), owing to no observable change in the POPC membrane micropolarity at different pHs (Fig S18-19).

**Figure 2:**
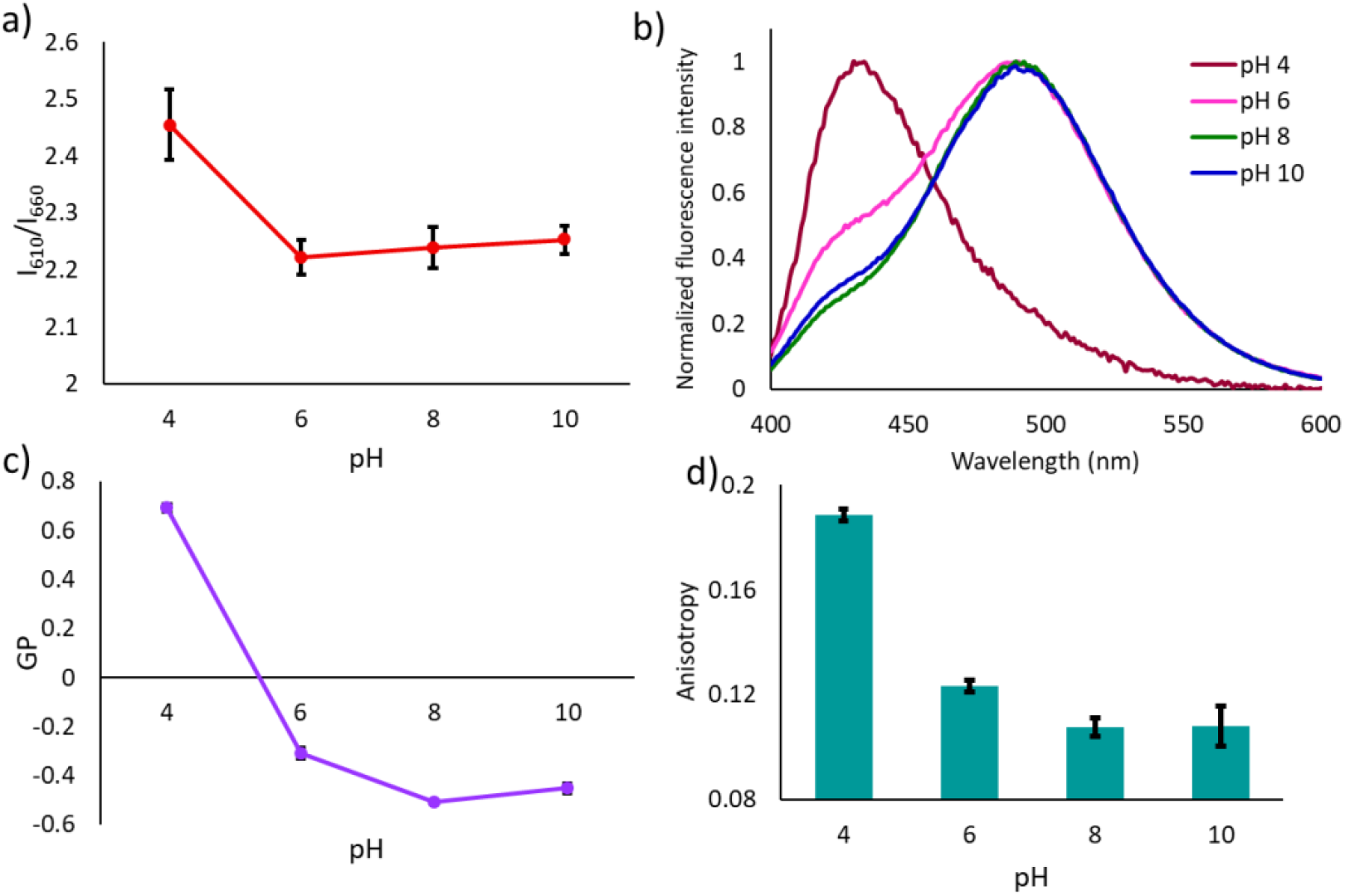
a) The scatter plot shows nile red I_610_/I_660_ ratio of the DDP suspension over varying pH at 45°C, N = 4, error bar = SD. b) Normalized emission spectra of laurdan in DDP suspension over varying pH at 45°C, N = 4, error bar = SD. c) The scatter plot shows laurdan GP values of the DDP suspension over multiple pH at 45°C, N = 4, error bar = SD. d) Laurdan anisotropy values of the DDP over varying pH at 45°C, N = 4, error bar = SD.

Next, laurdan was employed to investigate the order of DDP membranes at different pHs. Laurdan generalized polarization (GP) value provides a quantitative measure of the mobility of amphiphiles within a leaflet ^[31]^. Upon incorporation into membrane containing water accessible liquid-disordered phase, laurdan shows an emission maximum near 500 nm. In contrast, when incorporated into a solid-ordered phase, laurdan shows an emission maximum near 430 nm due to the reduced accessibility of water ^[27,36]^. Upon incorporation into DDP membranes at pH 4, laurdan emission maximum was found to be at 430 nm with a GP value of 0.69±0.01, alluding towards highly ordered membrane due to tightly packed DDP molecules (Fig 2b and 2c). At pH 6, laurdan showed two peaks, one at 430 nm corresponding to the ordered phase and the second one at 490 nm corresponding to the liquid-disordered phase with a GP value of −0.3±0.01. Upon increasing the pH to 8 and 10, the peak intensity corresponding to the ordered phase (at 430 nm) decreased, with GP values of −0.5±0.08 and −0.45±0.02, respectively. This shows that upon increasing the pH, the order (packing) of the DDP membrane decreases. The decrease in membrane order with increasing pH corroborates with increased membrane micropolarity.

To confirm that this change in membrane order with varying pH is unique to DDP, we measured the GP of two different PL membrane systems at different pHs, including POPC (melting temperature, −19°C) and DPPC (melting temperature, 42°C) (see Fig. S1), which form highly disordered and ordered membranes, respectively, at room temperature (25°C). We observed that in the case of both the PL membranes, the GP value remained unchanged at different pHs, indicating no change in the membrane order over the course of the pHs studied (Fig S20). To characterize DDP membrane fluidity as a function of pH, we then evaluated the anisotropy of laurdan. An increase in membrane fluidity will allow free tumbling of laurdan in the membrane, which would be reflected as a decrease in its anisotropy ^[37]^. At pH 4, laurdan anisotropy in the DDP membrane was found to be 0.18±0.02 (Fig 1d). Upon increasing the pH to 6 and 8, the anisotropy decreased to 0.12±0.02 and 0.10±0.02, respectively, after which it remained unchanged (Fig 2d). Therefore, with increasing pH, DDP membrane fluidity increased as one would expect based on the decrease observed in the GP values. This feature of the DDP membrane further highlights the tuneable nature of these membranes as a function of pH, and this property stems from the different protonation states. This pH-responsive changes in the DDP membrane’s properties underscores its adaptive nature to different environments. This has a potential functional aspect as membrane permeability is directly proportional to membrane lipid packing. Therefore, modulation in membrane properties in response to a change in pH would regulate membrane permeability thereby, the capacity for exchange of matter across the membrane boundary.

### 2c. CBC (Critical Bilayer Concentration) estimation of DDP membranes at different pHs

To further investigate the pH-dependent self-assembly process of DDP, its CBC was determined using NR and pyrene at pH 4 and 8. The NR I_610_/I_660_ ratio was plotted as a function of DDP concentration. As mentioned earlier, NR emission undergoes a blue-shift upon partitioning into hydrophobic regions. Thus, an increase in the I_610_/I_660_ ratio reflects the formation of self-assembled structures ^[34]^. A steady value of this ratio that continues to remain unaltered even when there is a further increase in DDP concentration, indicates stable membrane formation. A sharp increase in the I_610_/I_660_ ratio was observed in pH 8 at 0.1 mM DDP concentration (Fig 3a and Fig. S21). At pH 8, blue-shift of the emission light and the I_610_/I_660_ ratio reached a steady value at a relatively lower concentration when compared to pH 4 (Fig. S21-S22). In all, the CBC at pH 4 and 8 were found to be 0.9±0.1 and 0.7±0.1 mM, respectively, indicating that DDP assembles more readily at pH 8 (Fig 3a). To compare the CBC of pure DDP membrane with a FA system, we determined the CBC of lauric acid, which is a FA with the same aliphatic chain length as DDP (C12; Fig. S1). The CBC of lauric acid was found to be much higher at ~6 mM at pH 8 (Fig. S23), highlighting the influence of the phosphate head-group on the self-assembly process. Possessing such a low CBC as DDP is of extreme advantage in the prebiotic context as the presence of high concentrations of solutes in a dilute prebiotic pool would have been challenging, which poses difficulty for compartment formation ^[3–5]^.

**Figure 3:**
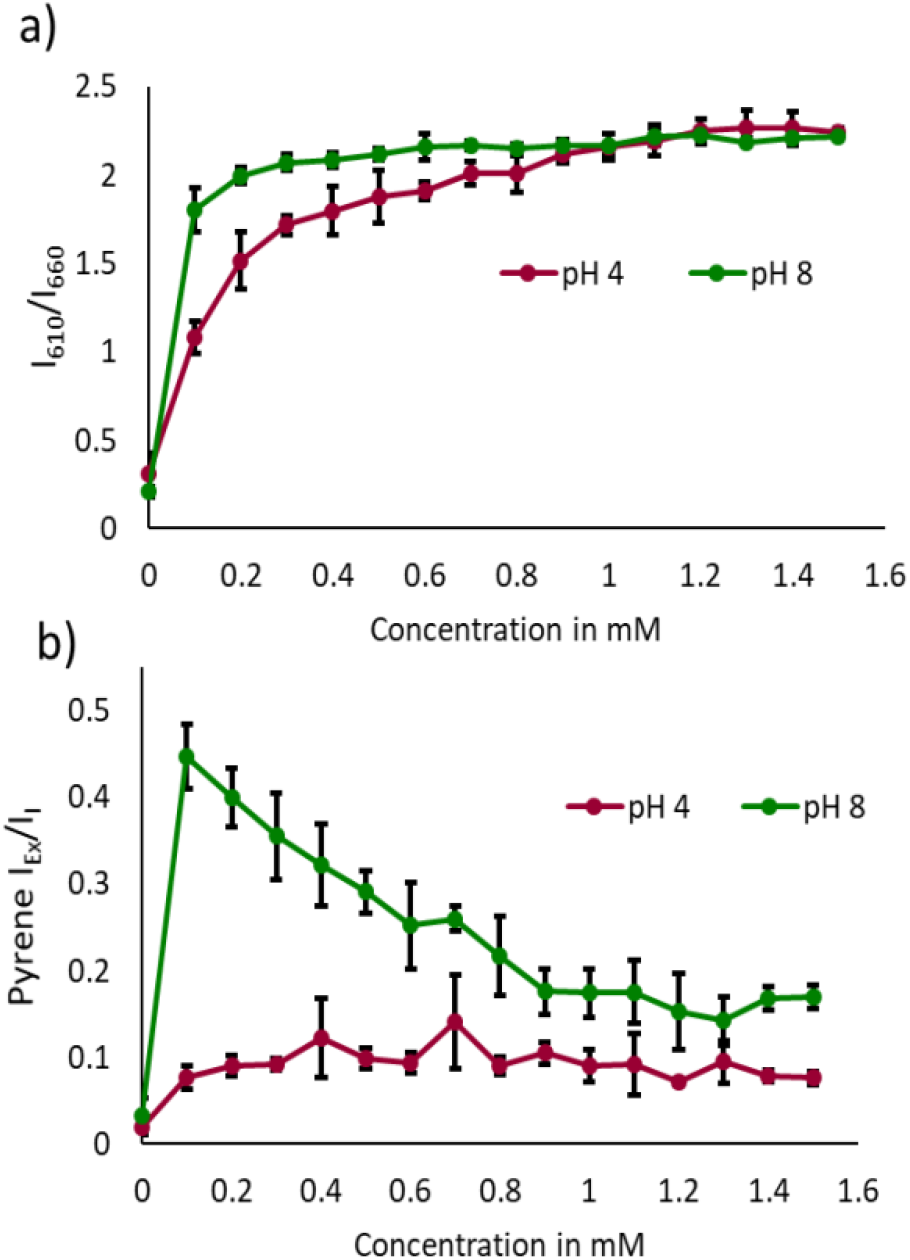
a) The scatter plot shows nile red I_610_/I_660_ ratio of DDP suspension at pH 4 and 8, plotted as the function of DDP concentration. N = 3, error bar = SD. b) Pyrene I_Ex_/I_I_ ratio of DDP suspension at pH 4 and 8, plotted as the function of DDP concentration., N = 3, error bar = SD.

Next, we plotted the pyrene I_1_/I_3_ ratio as a function of DDP concentration to crosscheck the CBC values obtained with NR ^[34]^. We observed a sharp decrease in the I_1_/I_3_ ratio with increasing DDP concentration, which reached a steady value similar to the NR I_610_/I_660_ ratio, indicating the formation of stable vesicles. The CBC at pH 4 and 8 were found to be 1±0.1 and 0.9±0.1 mM, respectively, when using the pyrene I_1_/I_3_ ratio (Fig S24-S25). In addition to the five monomer vibronic peaks between 370 to 400 nm, pyrene also shows a broad emission peak near 460 to 500 nm because of excimer formation ^[26,27]^. Pyrene excimerization is considered to occur due to the formation of excited-state dimer by the interaction between the ground-state and excited-state pyrene monomers owing to their close proximity ^[27,38]^. Thus, excimerization points toward the self-assembly process itself and the nature of aggregation too. To quantify the excimer formation, the intensity ratio of 470 nm (I_Ex_) and 372 nm (I_1_) was plotted as a function of DDP concentration. In pH 8, high intensity emission corresponding to excimerization was observed at 0.1 mM DDP, which gradually decreased with increasing DDP concentration (Fig 3b). Literature suggests that excimer formation at low surfactant concentration reflects premicellar aggregation ^[39,40]^. The continuous decrease in I_1_/I_3_ ratio at such low DDP concentration suggested the progressive increase in the self-assembled structures. With increasing DDP concentration, pyrene excimerization decreased as pyrene molecules get distributed in the DDP assemblies and the probability to find more than one pyrene in an assembly decreased at pH 8 ^[40]^. However, at pH 4, pyrene excimerization was found to be absent and unaffected by changing DDP concentration. The absence of a strong excimer emission peak at low (below CBC) DDP concentration at pH 4, suggests that pyrene monomers are unable to form excimers inside the aggregates (Fig 3b). Models suggest that pyrene excimerization is diffusion-limited in 2D space ^[38]^. The 2D model of diffusion-controlled pyrene excimerization was also validated for pyrene labelled phospholipid membrane (py10-PC/POPC and py10-PC/DMPC) and with free pyrene incorporated in DMPC membrane ^[41]^. Therefore, we hypothesize that the low I_Ex_/I_1_ ratio in pH 4 over varying DDP concentrations could reflect the limited diffusion of pyrene monomers in the DDP membranes. Additionally, higher anisotropy and GP value of DDP membrane at pH 4, reflect the rigid nature of the membrane that could prevent the diffusion of pyrene monomers.

To confirm the aforesaid, the crystalline nature of the DDP membranes at pH 4 was investigated by analysing the dried film of a DDP membrane suspension using Powder X-ray Diffraction (PXRD). Multiple sharp peaks were observed in the XRD pattern at pH 4 (Fig S26). Such peaks and patterns were absent in the XRD pattern of DDP suspension at pH 8. The appearance of sharp multiple peaks in periodic order at pH 4, denoted the crystalline nature of the membrane ^[42]^. To confirm that the sharp peaks at pH 4 were indeed coming from the DDP membrane, we extracted the DDP molecules from the pH 4 vesicle suspension in chloroform:methanol::2:1 and analysed its dried film XRD pattern. The absence of any peaks in the extracted samples confirmed that the sharp XRD peaks were indeed a characteristic of DDP membranes at pH 4 under aqueous conditions as DDP cannot assemble into vesicles in chloroform:methanol::2:1.

## 3. Effect of dodecanol (DOH) on the self-assembly and properties of DDP membrane

The addition of fatty alcohols has been shown to stabilize FA vesicles in alkaline pH and increase membrane thermostability ^[8,15,24]^. Contrarily, fatty alcohols are also known to induce FA vesicle destabilization and oil-droplet formation when added in a high molar ratio ^[8,15]^. Given these observations, we aimed to discern the effect of the presence of fatty alcohols when added in conjunction with DDP to result in membranes. Towards this, we mixed DOH with DDP to prepare DDP-DOH mixed membrane systems in different molar ratios i.e. DDP:DOH::1:1, DDP:DOH::2:1 and DDP:DOH::4:1. The turbidity measurements indicated the formation of higher-order aggregates in all three mixed systems over varying pH. Unlike in the pure DDP system, low turbidity of DDP:DOH::1:1 and DDP:DOH::2:1 mixed systems at pH 2, indicated the absence of large crystalline aggregates in the respective suspensions (Fig. S27-S28). Upon further decreasing the ratio of DOH to DDP in DDP:DOH::4:1, the turbidity at pH 2 increased, similar to what was observed in the pure DDP system (Fig S29). However, the high turbidity of the three mixed systems at alkaline pH (pH 10), when compared to the pure DDP system, alluded to an increase in vesicle formation in the presence of DOH (Fig 4a). Further, the pyrene I_1_/I_3_ ratio for all the three mixed systems mentioned above, across varying pH, indicated the formation of mixed aggregates with hydrophobic regions (Fig S30-S32). The I_1_/I_3_ ratio of the three mixed systems was significantly lower at pH 10 when compared to the pure DDP system, suggesting better shielding of pyrene molecules from the bulk water (Fig S33). Under the microscope, vesicles were observed from pH 3 to 10 in both DDP:DOH::1:1 and DDP:DOH::2:1systems (Fig. 4b and Fig. S34). However, upon decreasing the DOH molar ratio in DDP:DOH::4:1 mixed system, vesicles were observed from pH 2 to 10 similar to what was seen in the only DDP system (Fig. 4b and Fig. S34). Therefore, the presence of DOH at high concentrations decreased the solubility of DDP molecules in the aqueous phase and thus hamper vesicle formation at pH 2. However, at alkaline pH, DOH can act as H-bond donor and potentially H-bond with the negatively charged phosphate group of DDP, thereby, stabilizing vesicles as is reflected by an increase in the turbidity values (Fig 4a). The average size of the vesicles present in the three mixed systems at pH 10 was found to be significantly higher than the pure DDP system at pH 10 (Fig 4c). This increased vesicle size and turbidity, along with a decrease in pyrene I_1_/I_3_ ratio, highlights the vesicle stabilizing ability of DOH at alkaline pH, potentially by acting as H-bond donor.

**Figure 4:**
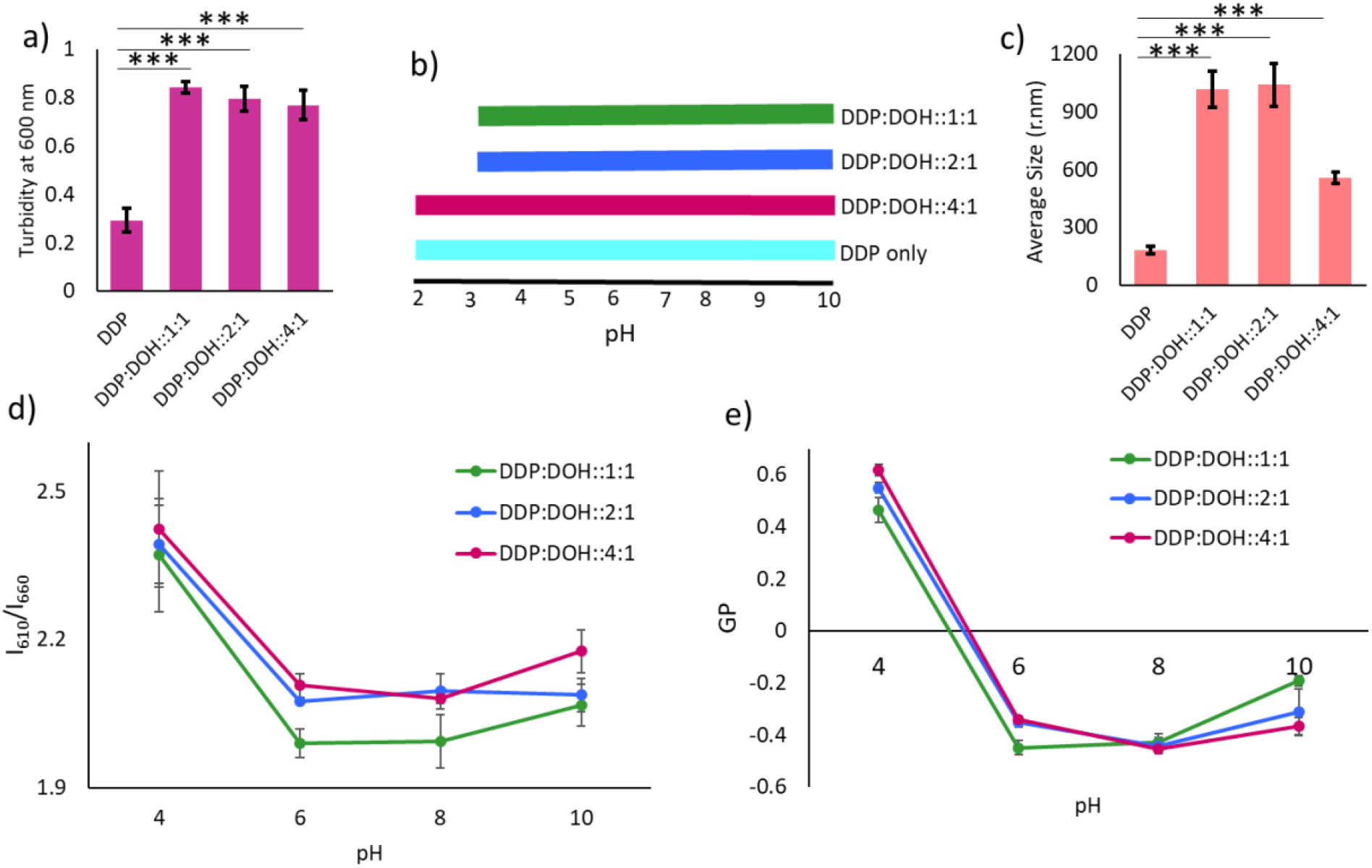
a) Turbidity measurements at 600 nm of 10 mM of DDP and different DDP:DOH mixed membrane systems at pH 10 recorded at 45°C, N = 3, error bar = SD. The comparison between the only DDP with the three DDP-DOH mixed membrane systems showed a difference that was found to be significant with a p-value of < 0.001 based on a two-tailed t-test. b) Shows the membrane formation ability of DDP and different DDP:DOH mixed systems over varying pH. The ability of a system to form vesicles over the range of pH is represented by the colored horizontal bars. c) Average size (nm) of the population of DDP and different DDP:DOH mixed systems at pH 10 recorded at 45°C. N = 4, error bar = SD. The comparison between the only DDP with the three DDP-DOH mixed membrane systems showed a difference that was found to be significant with a p-value of < 0.001 based on a two-tailed t-test. d) Shows nile red I_610_/I_660_ ratio of the DDP and different DDP:DOH mixed systems over varying pH at 45°C, N = 4, error bar = SD. e) Shows laurdan GP values of the DDP and different DDP:DOH mixed systems over varying pH at 45°C, N = 4, error bar = SD.

To investigate the physicochemical properties of the DDP:DOH mixed membranes, we used NR and laurdan probes. For all the three mixed systems, i.e. DDP:DOH in 1:1, 2:1 and 4:1 ratios, the change in micropolarity and GP values were evaluated as a function of pH. The trend of change in NR I_610_/I_660_ ratio with a change in pH was similar to what was seen in the pure DDP membrane system. The I_610_/I_660_ ratio was high in pH 4 and it decreased with an increase in pH indicating the increase in membrane micropolarity (Fig 4d). However, the I_610_/I_660_ ratio in all the mixed systems was overall lower when compared to the pure DDP membrane system (Fig. S35). The I_610_/I_660_ ratio was found to be lowest in the DDP:DOH::1:1 system, indicating high micropolarity (Fig 4d and Fig. S35). This suggested that the presence of DOH increased the membrane water accessibility. The GP values of all three mixed membrane systems decreased with an increase in pH owing to a decrease in membrane packing as observed in the case of only DDP membrane (Fig 4e). The GP values of the mixed systems were overall lower in comparison to the pure DDP membrane system, suggesting a decrease in membrane packing in presence of DOH (Fig 4e and Fig. S36). As the melting temperature of DOH (24°C; see Fig. S1) is much lower than that of DDP (42-45°C; see Fig. S1), and the GP values were recorded at 45°C, the presence of DOH potentially impedes DDP membrane packing and increases water accessibility. Similar to the pure DDP membrane, the decrease in laurdan anisotropy showed a decrease in the fluidity of all the mixed membranes upon an increase in pH. The fluidity was highest in pH 4 (with anisotropy value of 0.18) and decreased till pH 8 (with an anisotropy value of 0.1), after which it remained unaltered (Fig. S37-S39).

## 4. High-temperature behaviour of DDP and DDP:DOH mixed membranes at different pH

To understand the temperature-dependent phase behaviour of DDP-based membranes, the GP values were measured from 35°C to 65°C at pH 4 and 8 and were compared with values obtained for LA and myristoleic acid (MOA) membranes at pH 8. For the pure DDP membranes at pH 4, the GP value was 0.71±0.05 at 35°C corresponding to a highly ordered phase. With increasing temperature, it decreased owing to increased thermal motion (Fig. 5a and Fig. S40). At 65°C, it decreased to 0.48±0.01 indicating a decrease in membrane packing. For the DDP membranes at pH 8, the GP value at 35°C was −0.32±0.01 reflecting the disordered nature of the membrane (Fig. 5a). With increasing temperature, the GP value further decreased to −0.62±0.05 at 65°C, indicating a highly disordered DDP membrane (Fig. 5a). This highlighted the thermostability of the DDP membrane, especially at low pH (pH 4) against high temperature-induced membrane dissolution. To understand the effect of DOH on this, the GP of the DDP:DOH::1:1 mixed system was also monitored from 35°C to 65°C as the presence of fatty alcohol is shown to increase FA membrane thermostability ^[24]^. As shown in Fig 5a, in DDP:DOH::1:1 mixed system, the GP decreased from 0.58 to 0.15 upon increasing the temperature from 35°C to 65°C, at pH 4 (Fig. S40). At pH 8, the GP decreased from 0.18 to −0.61 upon increasing the temperature from 35°C to 65°C for the DDP:DOH::1:1 mixed system, which reflects a drastic decrease in the membrane order (Fig 5a). This shows that DOH can have a dual effect on membrane packing at pH 8; at low temperature, DOH increased the membrane packing, while at high temperature it decreased the membrane packing. For the LA, the GP was found to decrease from −0.30±0.01 to - 0.72±0.02 moving from 35°C to 65°C (Fig 5a). Upon plotting the decrease in GP for all the systems, the highest degree of decrease (0.8±0.01) was found for DDP:DOH (1:1) system at pH 8 (Fig 5b). This was followed by DDP:DOH (1:1) system at pH 4, which showed a decrease of 0.43±0.006. For the pure DDP and LA membranes (both at pH 8), the decrease in GP was 0.29±0.007 and 0.42 ±0.01, respectively, when the temperature changed from 35°C to 65°C. This highlighted that DDP membranes are relatively more resistant to temperature-induced decrease in membrane packing than LA membranes (Fig 5b). The smallest change in GP (0.07±0.03) was observed for the MOA membrane as the membrane is loosely packed even at 35°C because of unsaturation in it (Fig. S1).

**Figure 5:**
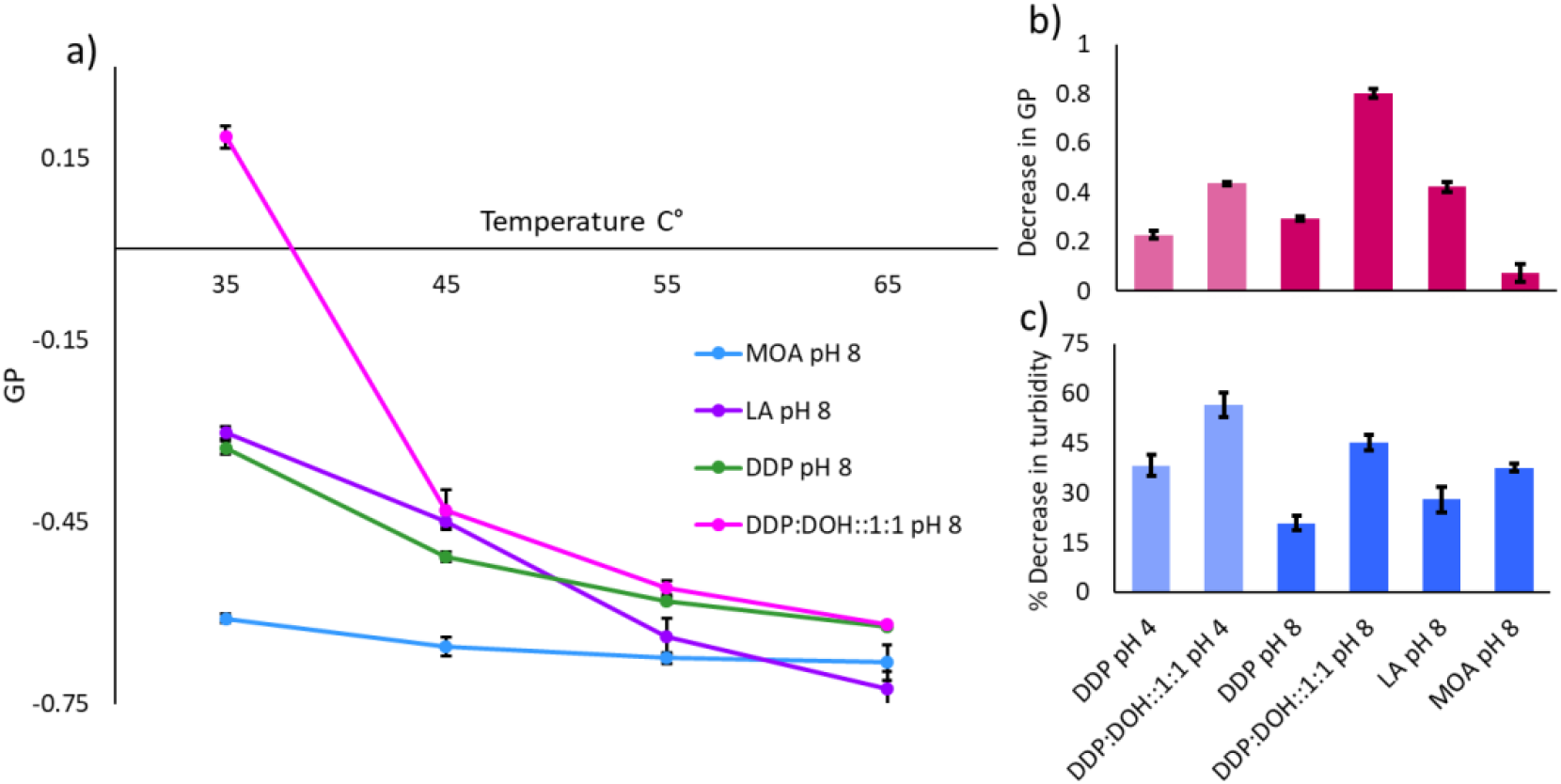
The graph shows laurdan GP values of the different membrane systems over varying temperatures at pH 8. N = 4, error bar = SD. b) The bar plot represents the decrease in laurdan GP values for different membrane systems upon increasing the temperature from 35°C to 65°C, at pH 4 and 8, respectively. N = 4, error bar = SD. c) The bar plot represents the decrease in the turbidity at 600 nm for different membrane systems upon increasing the temperature from 35°C to 65°C, at pH 4 and 8, respectively. N = 4, error bar = SD.

To understand the change in vesicle morphology at elevated temperatures, we next monitored the turbidity of the suspensions and plotted the change in turbidity upon increasing the temperature from 35°C to 65°C. In all the systems, a decrease in turbidity was observed upon increasing temperature (Fig. 5c). The highest percentage of decrease was observed for DDP:DOH::1:1 mixed system at pH 4 (56.7±3.6), followed by the DDP:DOH::1:1 system at pH 8 (45.2±2.3) (Fig 5c). The least decrease was observed for the pure DDP system at pH 8 (20.9±2.1). Using decanoic acid and decanoic acid-decanol (1:1 molar ratio) mixed membranes, studies have reported an increase in micellar and monomer population at the expense of vesicles, and also a decrease in vesicle lamellarity when heated above 60°C ^[32]^. As vesicle lamellarity and number are both known to linearly correlate with suspension turbidity, the decrease in turbidity observed here for all systems could therefore indicate a decrease in vesicle number and/or lamellarity ^[25]^. Since membrane order is inversely correlated with membrane permeability, a decrease in membrane packing at elevated temperature as observed here should facilitate the exchange of matter ^[24]^. At pH 8, DDP and DDP-DOH mixed membrane would possess high permeability similar to FAs membrane (LA). However at pH 4, because of the tight packing (high GP), DDP and DDP-DOH mixed membrane would possess extremely low permeability which can be counterbalanced by increasing the temperature.

## 5. Comparison of DDP membrane properties with different model protocellular membranes

In order to understand and correlate DDP membrane properties with other model protocellular membrane systems, we compared the micropolarity and order of DDP membranes with different PLs (1-palmitoyl-2-oleoyl-glycero-3-phosphocholine, POPC; 1,2-dipalmitoyl-sn-glycero-3-phosphocholine, DPPC; 1,2-dimyristoyl-sn-glycero-3-phosphocholine, DMPC; 1,2-di-dodecanoyl-sn-glycero-3-phosphocholine, DLPC; 1,2-di-decanoyl-sn-glycero-3-phosphocholine; DDPC) and FAs (Dodecanoic acid, LA; Decanoic acid, DA; Myristoleic acid, MOA; and Oleic acid, OA) membranes, of varying chain length and melting point (Fig. S1), both at pH 4 and 8 at 45°C. NR I_610_/I_660_ ratio and laurdan GP for four different phospholipids were determined at both pH 4 and 8. However, for all the four FA systems, it was determined at pH 8 as FAs don’t assemble into vesicles at pH 4. As shown in the Fig 6a, at pH 8, the NR I_610_/I_660_ ratio for all the four FA systems was between 1.50 to 1.68, indicating high membrane micropolarity. For all the four phospholipids, the I_610_/I_660_ ratio was between 1.86 to 2.16, indicating comparatively low membrane micropolarity (water accessibility). The I_610_/I_660_ ratio for DDP membranes at pH 8 was 2.2, suggesting that the DDP membranes possess similar polarity as that of phospholipid membranes at this pH (Fig. 6a). In terms of membrane order (GP) for FA membranes, the highest and lowest GP values were observed for LA (−0.41) and MOA (−0.73), respectively, at pH 8. For the DDP membranes at pH 8, the GP value was −0.50, which was similar to that of decanoic acid (C10, DA membranes) (−0.5) (Fig. 6a). Interestingly, though LA membranes possess the same chain length and similar melting point as that of DDP, DDP membranes are comparatively less packed at pH 8. For the phospholipid membranes at pH 8, DMPC had the highest GP (−0.26) and DDPC had the lowest GP (−0.49) at 45°C. The GP of DLPC (−0.40) was found to be similar to that of LA membranes (−0.41), which have the same aliphatic chain length. GP of DDPC (−0.5) was similar to that of DDP (−0.5), indicating the loose packing of the membrane. All the aforementioned data essentially highlighted the unique nature of the DDP membrane, where its micropolarity resembles a phospholipid membrane but its membrane packing is similar to that of both DA and DDPC membranes at pH 8. However, at pH 4, the I_610_/I_660_ ratio of the DDP membrane is 2.45, which is higher than the values obtained for the four phospholipid membranes used in this study (Fig S41). Importantly, the GP of the DDP membranes at pH 4 was much higher (0.69) than the phospholipid membranes studied here, reflecting its solid-ordered state, which was also reflected in the XRD pattern.

**Figure 6:**
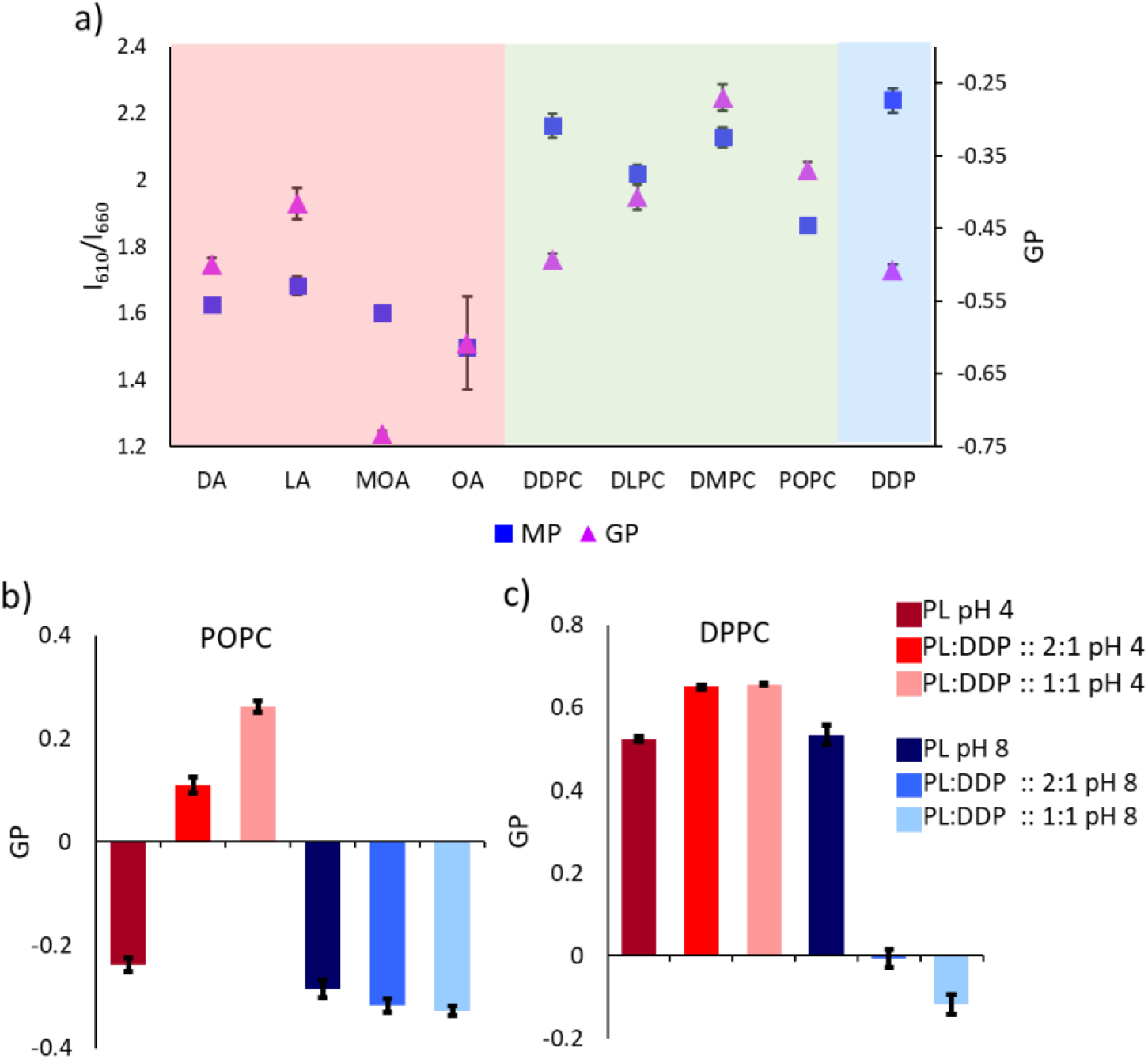
a) The scatter plot shows the nile red I_610_/I_660_ ratio (left y-axis) and laurdan GP values (right y-axis) of membranes i.e., different fatty acids, phospholipids and DDP as indicated on x-axis, at pH 8 and 45°C. N = 4, error bar = SD. b) The bar graph shows laurdan GP values of POPC and different POPC:DDP mixed membranes at pH 4 and 8, respectively, at 37°C. N = 4, error bar = SD. c) The bar graph shows laurdan GP values of DPPC and different DPPC:DDP mixed membranes at pH 4 and 8, at 37°C. N = 4, error bar = SD.

We next aimed to extend this unique tuneability to modulate the PL membranes by adding DDP to them. Recent studies reported an alteration in the membranes of multiple cancer cell lines resulting in increased membrane fluidity and decreased packing ^[43]^. In this context, the selective fusion of liposomes composed of PL to different cancer cell lines, exploiting the increased membrane fluidity of cancer cells, has also been reported ^[44–45]^. This highlights how the increase in the fluidity of the carrier liposome can lead to region-selective fusion. Therefore, to explore the modulation of membranes using DDP, we added DDP to different PL membranes in varying molar ratios. We monitored the change in the GP to evaluate the change in the packing of the mixed membrane in response to pH change at physiological temperature (37°C). As shown in Fig 6b, at pH 4 and 8, the GP of pure POPC membrane was −0.23±0.01 and −0.28±0.01 respectively, at 37°C. However, upon the addition of DDP to POPC (PL:DDP::2:1), the GP value changed to 0.11±0.01 and −0.31±0.01 at pH 4 and 8, respectively. Upon increasing the ratio of DDP (PL:DDP::1:1), the GP value further changed to 0.26±0.01 and −0.32±0.01 at pH 4 and 8, respectively (Fig. 6b). Similarly, in the case of DMPC, the addition of DDP at pH 4 led to an increase in membrane packing, while at pH 8 it led to a decrease in membrane packing (Fig. S42) as summarized in table ST2. We next studied DPPC, which has a high melting point (41 °C) and is also known to stay in ordered crystalline form at both pH 4 and 8 with a GP value of 0.53±0.02 for this analysis. The addition of DDP in 1:1 molar ratio resulted in a further increase in the order of the mixed DPPC membrane to 0.65±0.01 at pH 4. However, at pH 8, the GP decreased to −0.11±0.02 showing a drastic decrease in the membrane order (Fig 6c). Therefore, using three different PL membrane systems, with two ratios of DDP spiked in them, we demonstrated that the addition of DDP can lead to the tuneability of the resultant mixed PL membranes. At pH 4, it resulted in increased membrane packing while at pH 8 it led to a decrease in membrane packing owing to the protonation state of the DDP molecule (Table ST1). At pH 8, 19% of DDP stays di-deprotonated in the membrane resulting in repulsion between the doubly negatively charged headgroups, leading to loosening in the packing. Interestingly, the change in GP in response to pH when DDP is added to the milieu, is remarkably high.

## Conclusions

In this study, we characterized the pH-dependent self-assembly behaviour of DDP, an enticing candidate protocellular membrane. In contrast to FAs, which can assemble into vesicles only in a narrow regime of pH (pH 7-9), DDP alone can self-assemble into vesicles at a wide variety of pH, from pH 2 to 10. This highlights the compatibility of DDP vesicles with different prebiotically pertinent reactions that have been described to occur at a range of pH, including nonenzymatic oligomerization and replication reactions of the informational polymers occurring at a wide range. Systematic investigation of the properties of DDP membranes at varying pH revealed the markedly tuneable nature of these membrane systems. The micropolarity of DDP membranes was found to be lower at pH 4 and it increased as the surrounding pH increased. The packing and fluidity of these membranes were also found to be dependent on the surrounding pH, which decreased upon increasing pH. Such pH-responsive nature can aid in the functionalization of the compartment that, in turn, facilitates robust adaptation under different environmental conditions. Significantly, change in membrane micropolarity and packing in response to change in pH, would enable the modulation of the DDP membrane’s permeability. For e.g. at acidic pH (~pH 4), an increase in DDP membrane packing would decrease membrane permeability and protect the encapsulated material from dilution and molecular parasites. On contrary, in neutral to alkaline pH (~pH 8), the decrease in DDP membrane packing would increase the exchange of materials like building blocks, between the protocell and environment, affecting the rates of encapsulated reactions like templated RNA replication. Additionally, this pH responsiveness would also directly impinge on the interaction capability and binding efficiencies of different co-solutes, including nucleotide and amino acid monomers and oligomers.

DDP assemble readily into membranes with low CBC as compared to FAs. Interestingly, the CBC varies with varying pH. The addition of DOH facilitated vesicle formation at alkaline pH by potentially H-bonding with the deprotonated DDP molecules. Further, the presence of DOH decreased the overall packing of the mixed (DDP:DOH) membranes. On similar lines, DOH also increased the overall micropolarity of the mixed membranes. DDP and DDP:DOH::1:1 membranes were more stable at elevated temperatures when compared to FA membranes. Upon increasing the temperature, the DDP membrane packing decreased, however, the extent of this decrease was dependent on the pH and also the presence of DOH. As indicated earlier, this decrease in order would increase membrane permeability to facilitated the exchange of matter between the protocellular compartment and its environment ^[24]^. The decrease in suspension turbidity at elevated temperatures indicated the possibility of decreased membrane lamellarity. Next, we compared DDP membrane properties with different FAs and PLs membranes at pH 4 and 8, DDP membranes were found to be loosely packed and as dynamic as DA (C10) membranes at pH 8. However, the micropolarity of the membrane at this pH was similar to that of PL membranes. Nonetheless, unlike phospholipid membranes, the lower order in DDP membranes at neutral to alkaline pH, highlights its ability to support the emergence and sustenance of protocellular life forms.

Finally, we extended the pH-dependent tuneable nature of DDP to phospholipid membranes by doping the phospholipid membranes with DDP. The addition of DDP allowed for controlling the membrane order of the PL:DDP mixed membranes when subjected to changing pH. In few previous studies, PL:FA mixed membrane properties have been studied in the context of their role as model protocellular membranes ^[46]^. We propose that DDP-phospholipid mixed membrane systems would have been even more ideal in this context, due to being a malleable and robust class of prebiotically plausible pH-responsive model protocellular membranes. This pH-responsive nature of PL:DDP mixed membranes, not only underscores the implications of these systems in the context of protocell formation, but would also be relevant for the synthetic cell engineering purposes. Pertinently, RNA oligomers have also been shown to bind more efficiently with ordered crystalline lipid membranes ^[47]^ than disordered ones. The ordered nature of the DDP membranes in acidic pH could also help in the localization and concentration of RNA molecules. Moreover, the addition of DDP in PL-membranes provides an additional component for creating adaptable artificial cells, where one could program versatile membrane functions that respond to a pH change. Finally, after undertaking more systematic research, the tuneable nature of the DDP membranes can also be implemented for constructing pH-responsive synthetic cells that would allow for a selective and regiospecific release of drug molecules.

## Acknowledgements

The authors wish to acknowledge the Microscopy facility at IISER Pune. We are grateful to Dr. Amitabha Chattopadhyay for the useful discussions.

## Author contributions

S.S. S.D and S.R. designed the experiments. S.D performed the DLS data analysis and PXRD. KL performed the Foamability, foam stability assay and membrane thermostability experiments. S.S. performed the rest of the experiments. S.S., S.D. and S.R. analyzed the data. S.S., S.D. and S.R. wrote the manuscript.

