## Supplementary Information for "pH-responsive self-assembled compartments as tuneable model protocellular membrane systems"

### Experimental Section:

#### Materials:

All single chain amphiphiles used in this study, namely, mono-N-Dodecyl phosphate (DDP), Dodecanoic acid (LA), 1-Dodecanol (DOH), Decanoic acid (DA), Myristoleic acid (MOA) and Oleic acid (OA) were procured from Sigma Aldrich (Bangalore, India) and used without further purification. All the phospholipids used in this study, 1-palmitoyl-2-oleoyl-glycero-3-phosphocholine (POPC), 1,2-dipalmitoyl-sn-glycero-3-phosphocholine (DPPC), 1,2-dimyristoyl-sn-glycero-3-phosphocholine (DMPC), 1,2-di-dodecyl-sn-glycero-3-phosphocholine (DLPC), 1,2-di-decanoyl-sn-glycero-3-phosphocholine (DDPC) were purchased from Avanti Polar Lipids, Inc (AL 35007, United States) and used without further purification. All other chemicals used in this study were purchased from Sigma Aldrich (Bangalore, India). For all the experiments, nanopure water (with resistivity 18 MΩ-cm) was used.

#### Methods:

#### Vesicle suspension preparation:

The lipid suspensions (DDP, DDP:DOH, FA and PL) were prepared by rehydrating the thin lipid dry film with a buffer of the desired pH. The lipid film was prepared by drying the desired volume of chloroform solution of the lipids under nitrogen gas flow. Any trace amount of chloroform was removed by keeping the thin film under vacuum for another six hours. After rehydration with buffer, the suspension was heated for 1 hour at 45°C and vortexed occasionally to suspend the thin film properly. For pH 2 and 3, 100 mM Glycine-HCl buffer, for pH 4 and 5 100 mM Na-acetate buffer, for pH 6, 7 and 8, 100 mM K-phosphate buffer, for pH 9 and 10, 100 mM Glycine-NaOH buffer was used, respectively, to rehydrate the lipid dry film.

#### Microscopy:

Lipid samples were observed under 40X magnification using a Differential Interference Contrast (DIC) microscope AxioImager Z1 (Carl Zeiss, Germany), (NA = 0.75) to observe the presence of various higher-order aggregates. Typically ~10 µL of the lipid suspension was spread on a glass slide, covered and sealed with a glass coverslip and observed immediately at room temperature (25°C).

**Dynamic light scattering (DLS)**

To characterize the size distribution of the self-assembled structures (vesicle), 5 mM lipid suspension (DDP and DDP:DOH) was subjected to DLS using a 633 nm red laser from Malvern instruments. In a typical reaction, 1 ml of the lipid suspension was taken in a polycarbonate cuvette and analysed by vesicle size estimation on a Zetasizer Nano ZS90, (Malvern Panalytical Ltd., Malvern, UK) at 45°C. The data reproducibility was checked by performing at least three independent measurements. The correlation function (g) was plotted over time and the data was fitted into 1$^{st}$ and 2$^{nd}$ order exponential decay graphs. The decay constants were calculated from the graphs. The size distribution was plotted as a function of % intensity of the scattered light. The average size of the vesicle population (Z) was also plotted for different pH.

**Turbidity estimation:**

The turbidity of the lipid suspension was measured by taking the optical density of the suspension at 600 nm. Samples were loaded into a quartz cuvette and the turbidity was measured at 45°C using a UV-1800 UV-Vis Spectrophotometer (Shimadzu Scientific Instruments Inc., Columbia, USA).

**Powder X-ray Diffraction**

10 mM of aqueous vesicle suspension of DDP at pH 4 and pH 8, was drop-casted repeatedly on a glass slide to make a thick film. It was then air-dried, followed by vacuum drying for 24 hours at room temperature. Data was recorded on the Bruker D8 Advance X-ray diffractometer for these samples from 2° to 80°. The parameters for the same were as follows: scan speed = 0.5 second/step, increment: 0.2, rotation: 15.

**Foamability and foam stability assay:**

Glass vials (of diameter 1.2 cm and height 5 cm) containing 1 mL of DDP suspension of different pH (pH 2, 4, 6, 8 and 10) were kept in a cardboard box and the box was shaken vigorously by hand for 2 mins to generate foam. To look at the foamability of each suspension, the foam volume was measured right after the foaming process. Foam stability was characterized by measuring the volume of foam volume after different time points (2, 4 and 8 hours).

**Steady-state fluorescence analysis using nile red, laurdan and pyrene:**

The self-assembly behavior and physicochemical properties of the membranes were evaluated using solvatochromic dyes. In a typical reaction, 2 µl of 200 µM dye solution was added to 80 µl of the sample, to reach a final concentration of 5 µM (to maintain the dye to amphiphile molar ratio at 1:2000). This was followed by incubation of the suspension for 20 minutes at 45°C with shaking at 1000 rpm to let the fluorophores incorporate into the hydrophobic environment. The fluorescence readouts were taken at 90° angle on a FluoroMax 4 (HORIBA JOBIN VYON fluorescence spectrophotometer) with 150 W CW Ozone-free xenon arc lamp. The excitation and emission slit width were kept at 2 nm. For laurdan anisotropy measurements, excitation and emission slit width were kept at 4 nm. Lamp intensity variations were checked and corrected if required. To avoid any contribution coming from the scattering of the self-assembled structures within the fluorescence spectrum, the emission spectrum of a control sample was obtained without any added dye and this value was subtracted from the spectrum of the test sample.

**Nile red:** After the addition of nile red and incubation for the requisite period, the test suspensions were excited at 530 nm, and the emission spectra were collected from 550 nm to 750 nm. The intensity ratio at 610 and 660 nm was used to calculate the $I_{610}/I_{660}$ ratio.

**Pyrene:** After the addition of pyrene and incubation, the test suspensions were excited at 335 nm, and the emission spectra were collected between 350 to 550 nm. The ratio of peak 1 ($I_1$) at 372 nm and peak 3 ($I_3$) at 383 nm was used to calculate the $I_1/I_3$ ratio. To discern if there was excimerization, the intensity ratio of 470 nm ($I_{Ex}$) and peak 1 at 372 nm was used.

**Laurdan:** In the case of laurdan, after the incubation period, the test suspensions were excited at 370 nm and the emission spectra were collected from 400 to 600 nm. The generalized polarization (GP) was calculated by using the following equation:

$$GP = \frac{(I430 - I500)}{(I430 + I500)}$$

where, I430 and I500 represent fluorescence intensity at 430 and 500 nm, respectively.

For the laurdan anisotropy measurements, after the addition of laurdan and the incubation period, the test suspensions were excited at 370 nm, and the emission intensity was collected at 450 nm. The anisotropy was calculated by using the following equation:

$$\text{Anisotropy} = \frac{(I_{\mathrm{VV}} - GI_{\mathrm{VH}})}{(I_{\mathrm{VV}} + 2GI_{\mathrm{VH}})}$$

Where, $I_{VV}$ denotes intensity with vertical excitation and vertical emission. $I_{VH}$ denotes intensity with vertical excitation and horizontal emission. The instrument calculated the correction factor (G) for each measurement.

**Statistical analysis:**

All statistical analysis was performed using Microsoft Excel 2016. Two-tailed t-test was used to check the significance of the difference between the values within a system and also to compare between values of particular time points across systems. Values were considered statistically significant for values with $P < 0.05$.

**Supplementary Figures:**

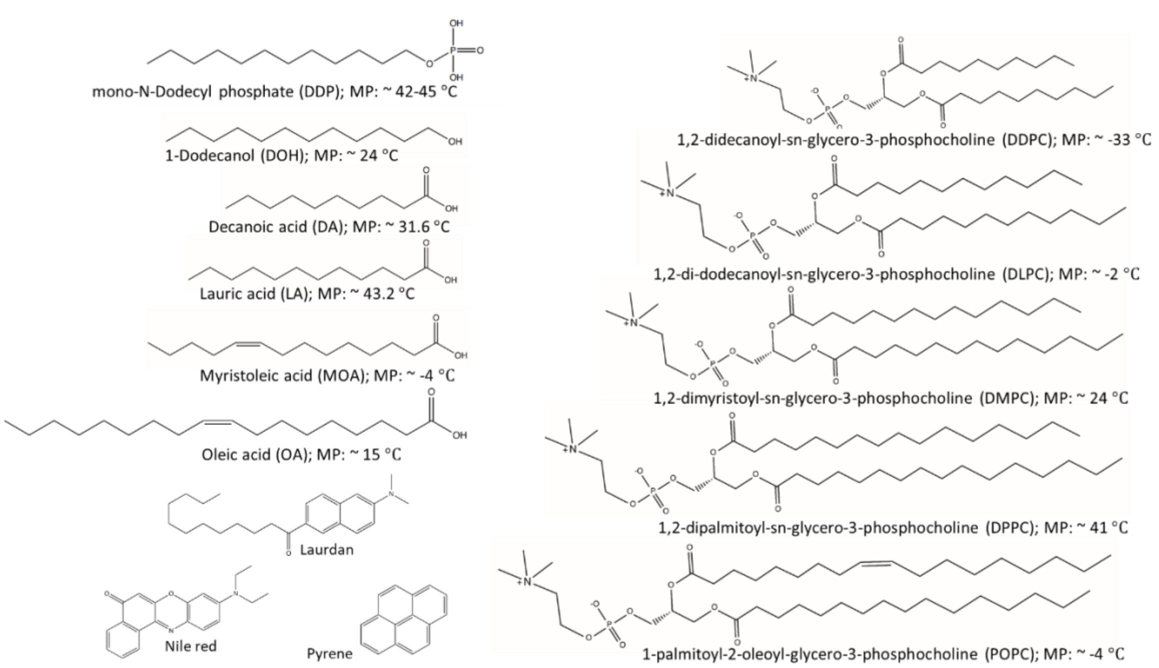


Figure S1: Chemical structures of the fatty acids (DA, LA, MOA and OA), DDP, DOH and phospholipids (DDPC, DLPC, DMPC, DPPC and POPC) along with their melting points (MP), and the fluorophores (laurdan, nile red and pyrene) used in this study. The structures were drawn using ChemDraw professional (PerkinElmer) 20.0. The melting temperatures were obtained from ChemSpider.

Pence, Harry E., and Antony Williams. "ChemSpider: an online chemical information resource." (2010): 1123-1124.

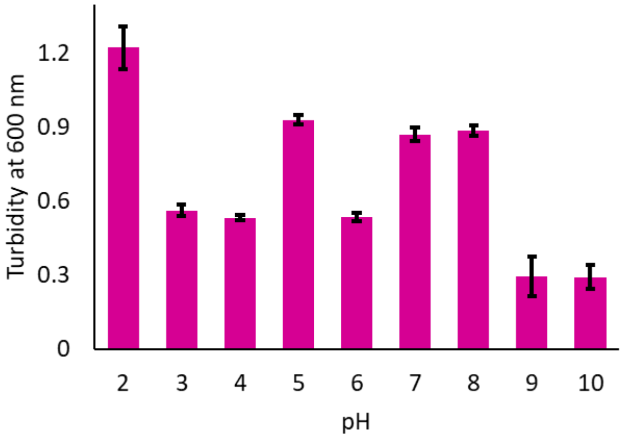


Figure S2: The bar plot shows the turbidity measurements obtained at 600 nm (y-axis) for the 10 mM DDP suspension at different pH (x-axis) and 45°C, N = 3, error bar = SD.

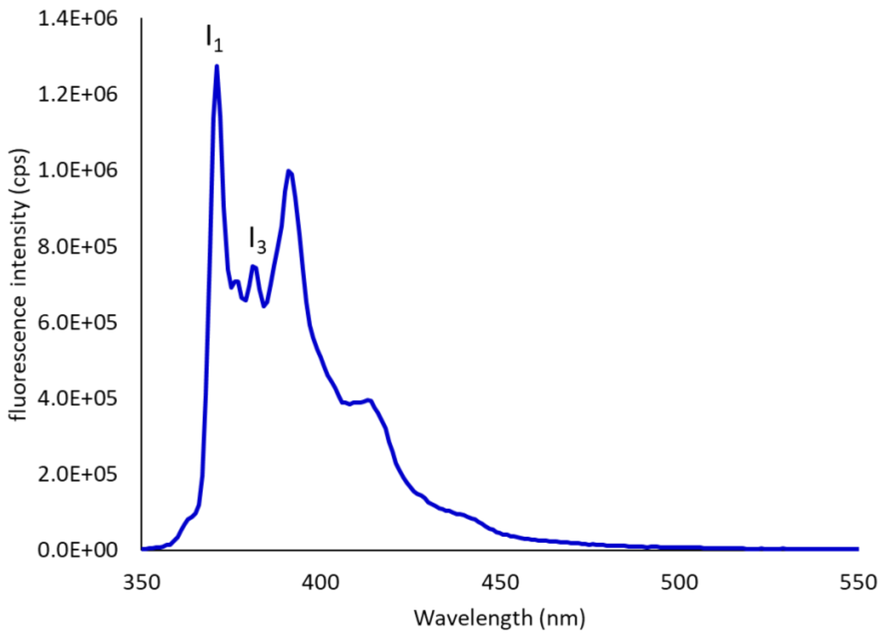


Figure S3: Emission spectrum of pyrene in water when excited at 335 nm. (N = 4). Five characteristic emission peaks were observed between 370 nm to 400 nm. The intensity ratio of $I_1/I_3$ indicate pyrene's microenvironment, where lower ratio denotes more hydrophobicity of the environment.

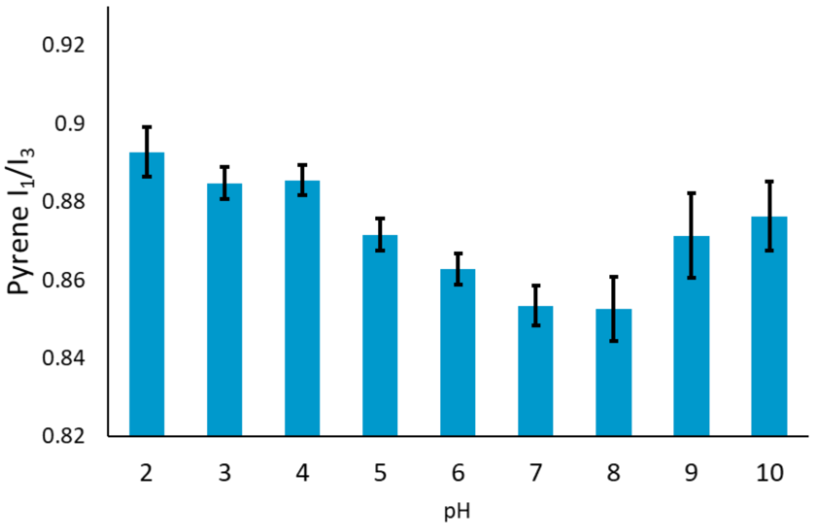


Figure S4: Pyrene $I_1/I_3$ ratio of 10 mM DDP suspension at different pH (x-axis) at 45°C, N = 4, error bar = SD. The $I_1/I_3$ ratio was found to be highest at pH 2, indicating less hydrophobicity of the environment.

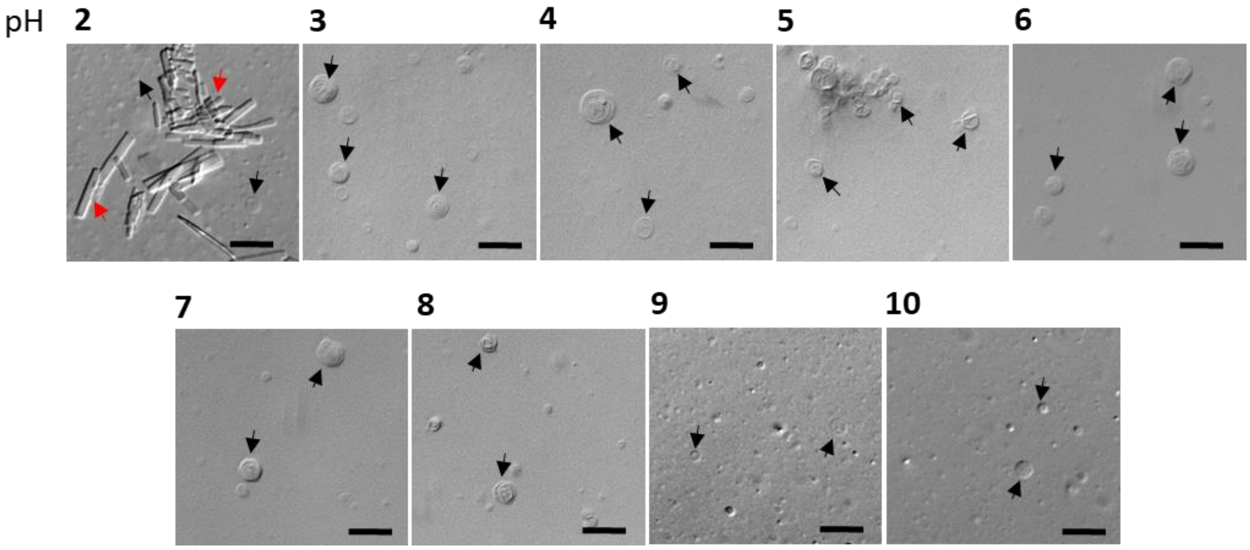


Figure S5: DIC microscopy images of DDP suspension at different pH. Black and red arrows indicate vesicles and crystalline aggregates, respectively. N = 3, Scale bar = 10 µm.

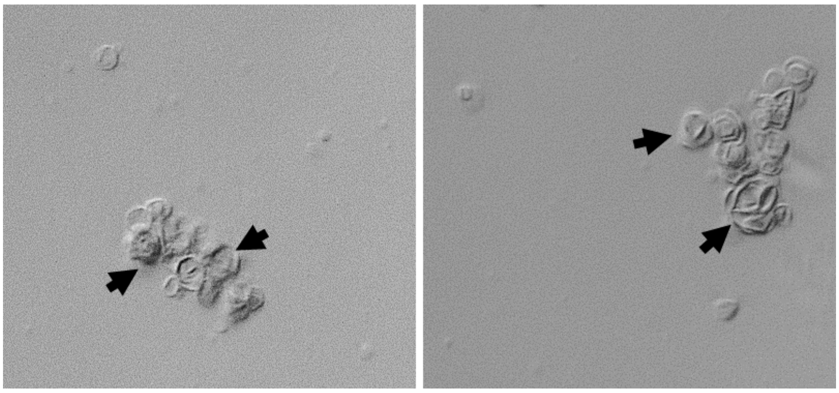

Figure S6: DIC microscopy images of DDP suspension at pH 8. Black arrows indicate clumped vesicles. N = 3, Scale bar = 10 µm.

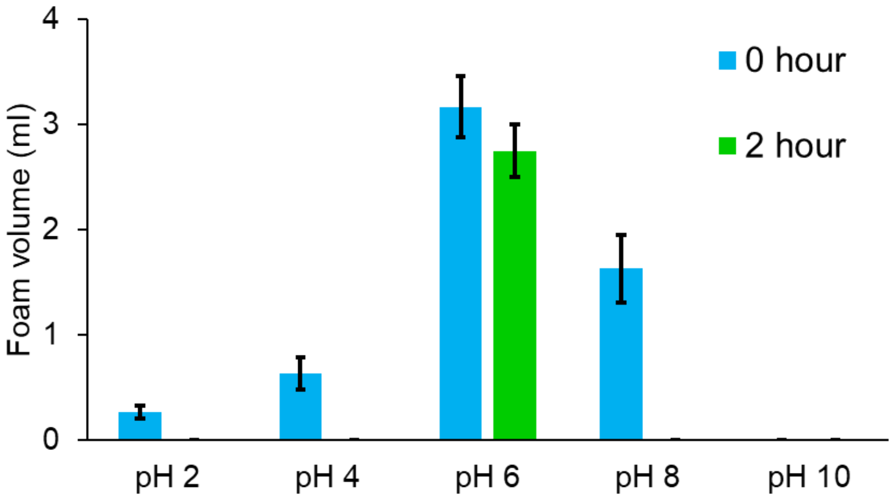


Figure S7: Foam stability assay of DDP suspensions. The bar graphs show the volume (ml) of foam formed in 5 mM DDP suspension (y-axis) at different pH (x-axis) at the initiation and after two hours of incubation at room temperature. N = 3, error bar = SD.

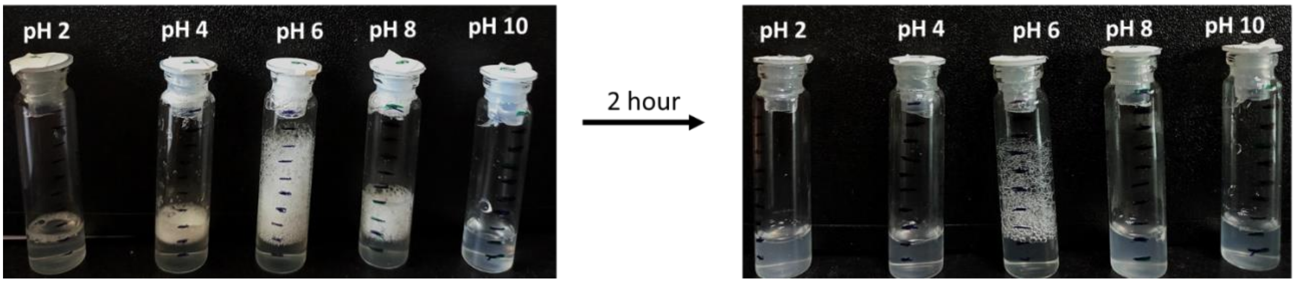


Figure S8: Images of glass tubes containing 5 mM DDP suspension at different pH after vigorous shaking. The images show the volume of foam (foamability) at the initiation and the volume remaining after two hours of incubation depicting foam stability at different pH. N = 3, error bar = SD.

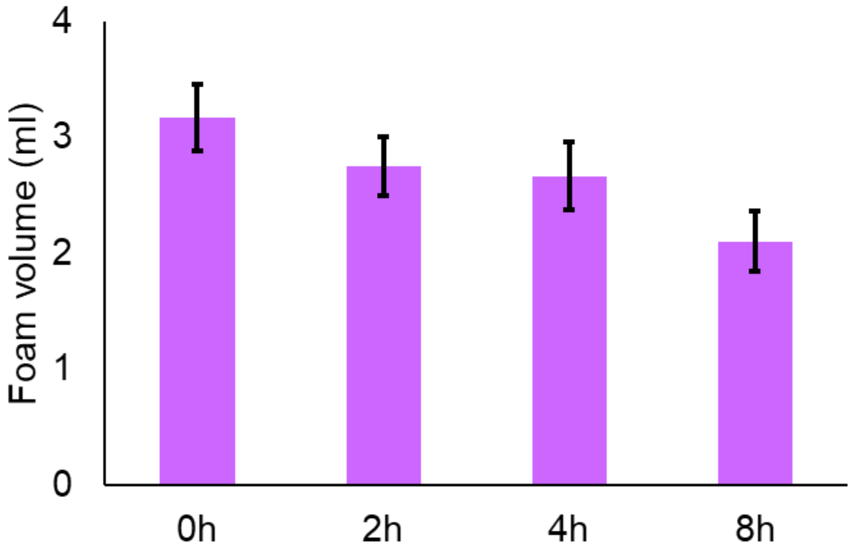


Figure S9: Foam stability assay of 5 mM DDP suspension at pH 6 after different incubation times (x-axis). The Y-axis shows the foam volume in ml. N = 3, error bar = SD.

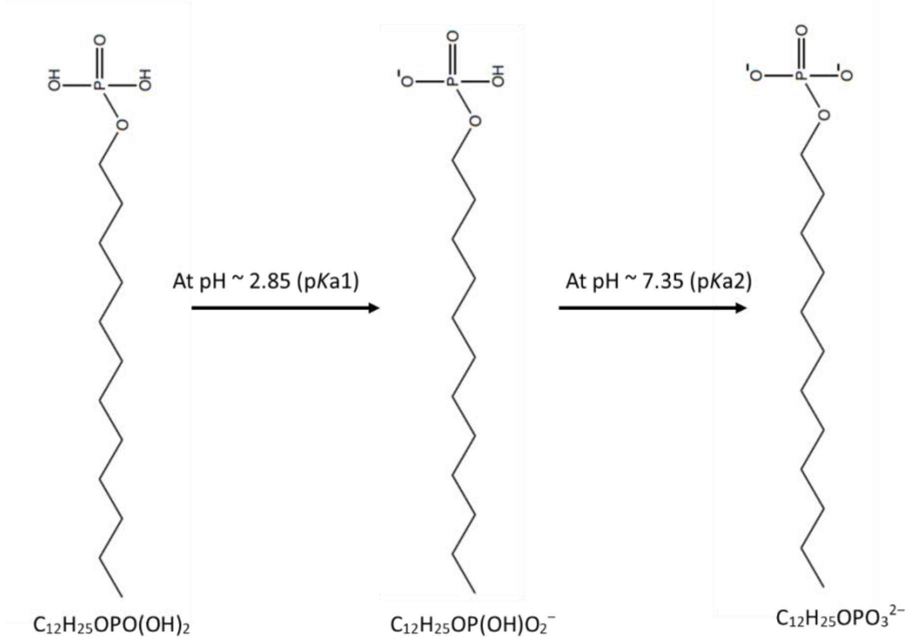


Figure S10: Molecular structural transition of DDP with respect to change in pH of the surrounding suspension. DDP possesses two known pKa; one at 2.85 (pKa1) and another at 7.35 (pKa2). At pH 2.85, 50% of the DDP population is in the monoprotonated state with the other half in fully protonated state. At pH 7.35, the monoprotonated state and doubly deprotonated state are in equal proportion.

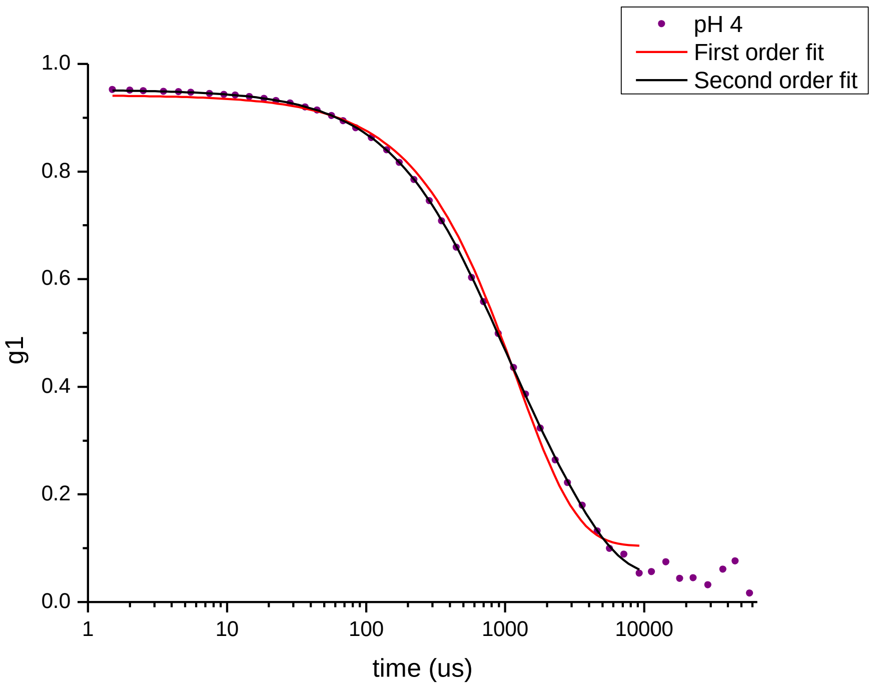


Figure S11: The DLS correlation function at pH 4. The DLS data was fitted using both first-order exponential decay curve (red) as well as second-order exponential decay curve (black). As observed, the correlation data perfectly fit the second-order exponential decay curve, indicating the heterogeneity in the size of self-assembled structures. N = 4.

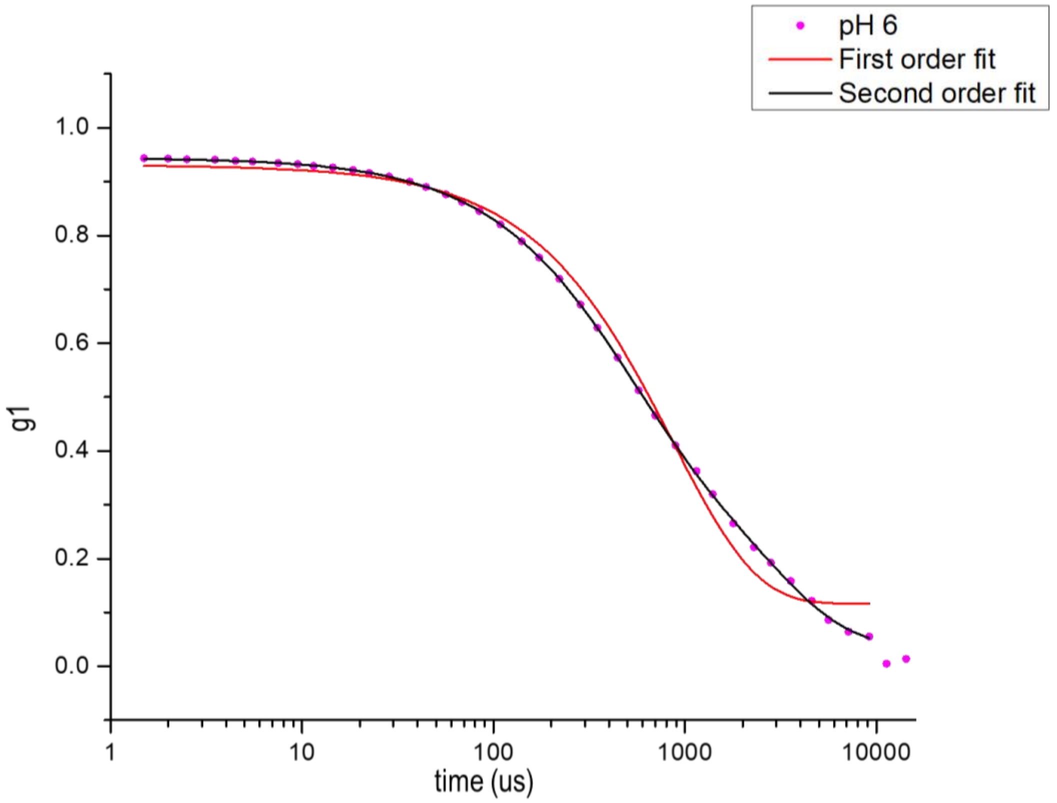

Figure S12: The DLS correlation function at pH 6. The DLS data was fitted by both first-order exponential decay curve (red) as well as second-order exponential decay curve (black). As observed, the correlation data perfectly fit the second-order exponential decay curve, indicating the heterogeneity in the size of self-assembled structures. N = 4.

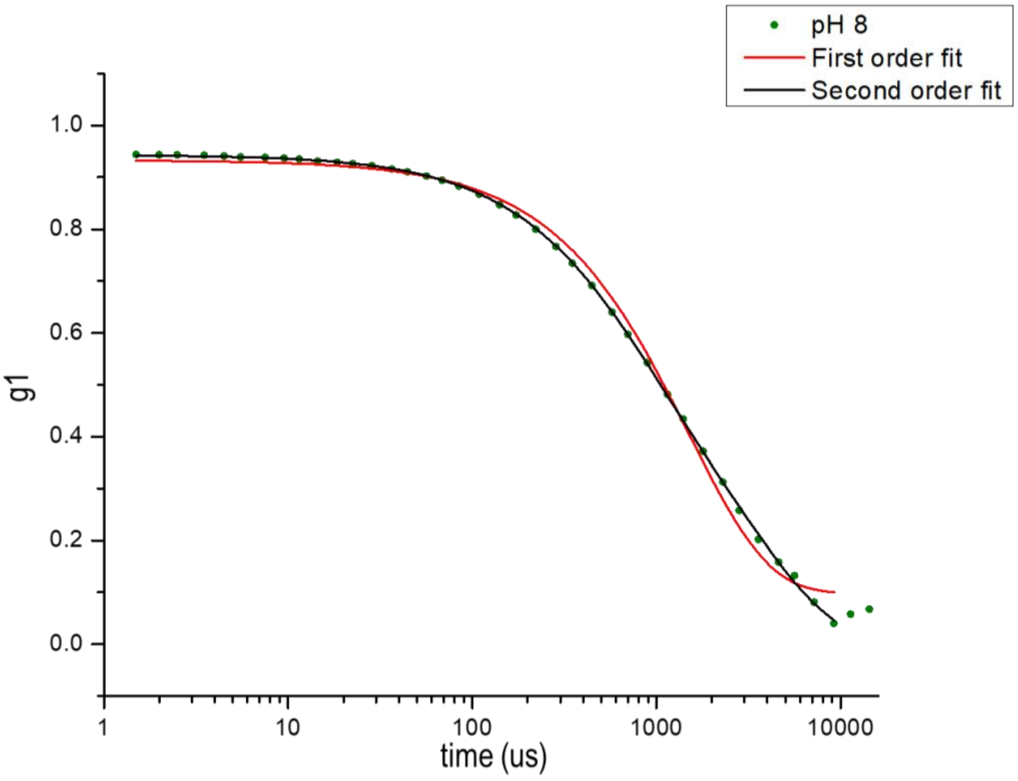


Figure S13: The DLS correlation function at pH 8. The DLS data was fitted by both the first-order exponential decay curve (red) as well as second-order exponential decay curve (black). As observed, the correlation data perfectly fit the second-order exponential decay curve, indicating the heterogeneity in the size of self-assembled structures. N = 4.

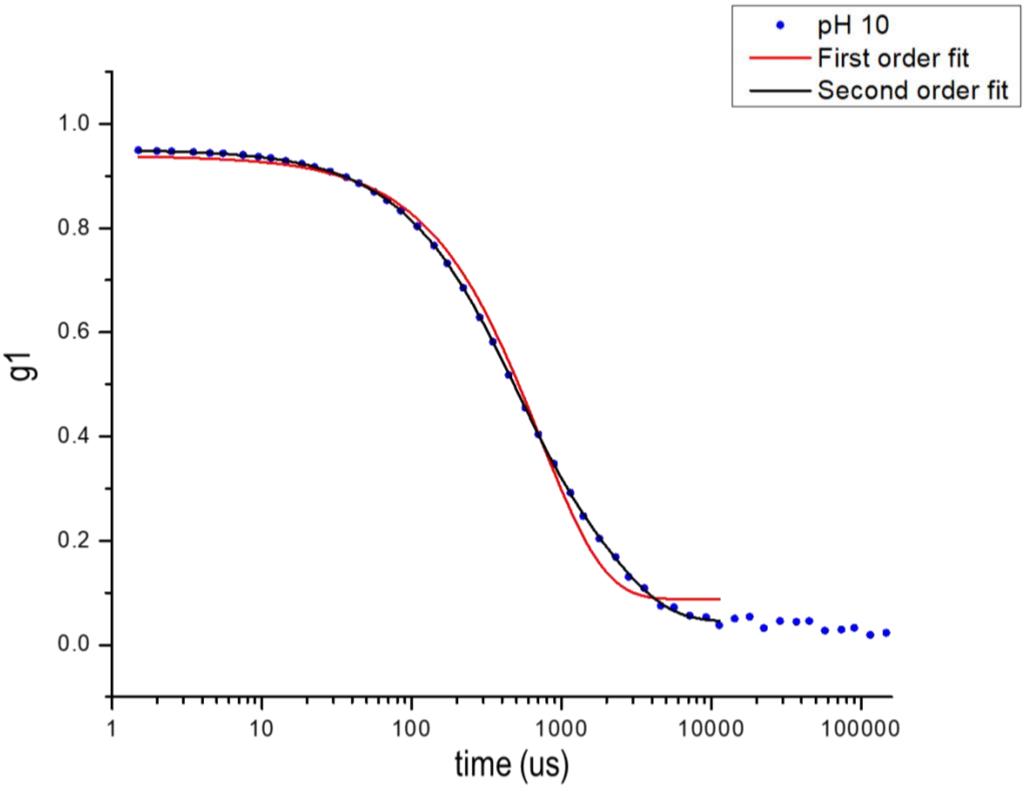


Figure S14: The DLS correlation function at pH 10. The DLS data was fitted by both first-order exponential decay curve (red) as well as second-order exponential decay curve (black). As observed, the correlation data perfectly fit the second-order exponential decay curve, indicating the heterogeneity in the size of self-assembled structures. N = 4.

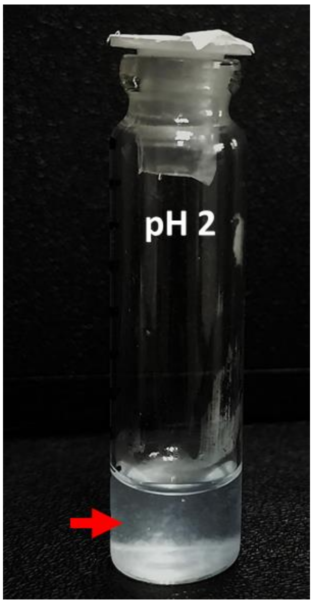


Figure S15: Image shows flocculation (precipitating aggregates) in 5mM DDP suspension at pH 2 after incubation (without shaking) for 5 minutes at 45°C. N = 4.

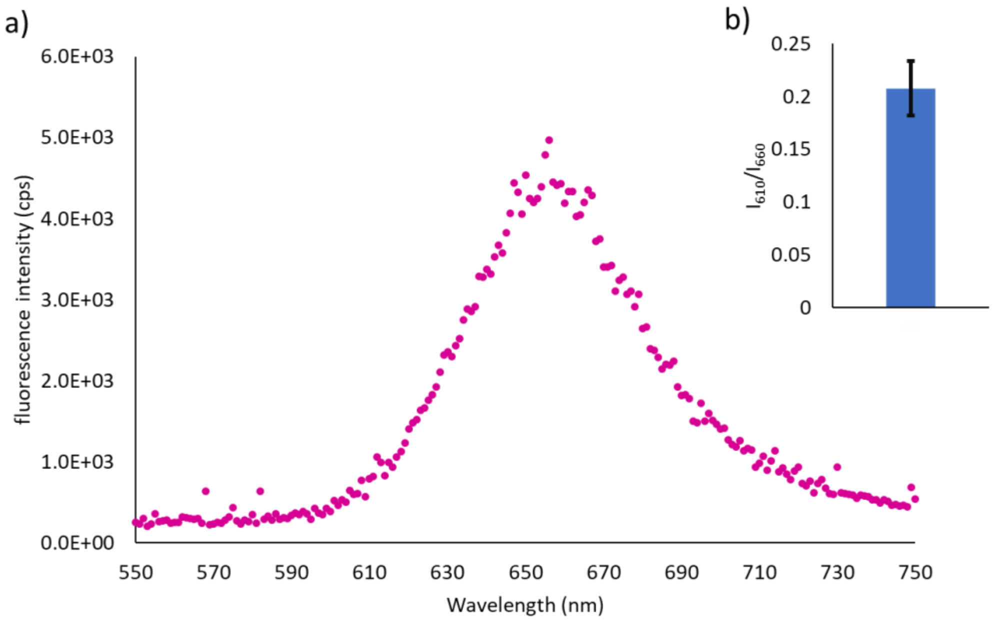


Figure S16: a) Emission spectrum of nile red in water when excited at 530 nm (N = 4). Emission maximum was observed at 660 nm. b) $I_{610}/I_{660}$ emission intensity ratio (0.2±0.02) of nile red in presence of water. N = 3, error bar = SD.

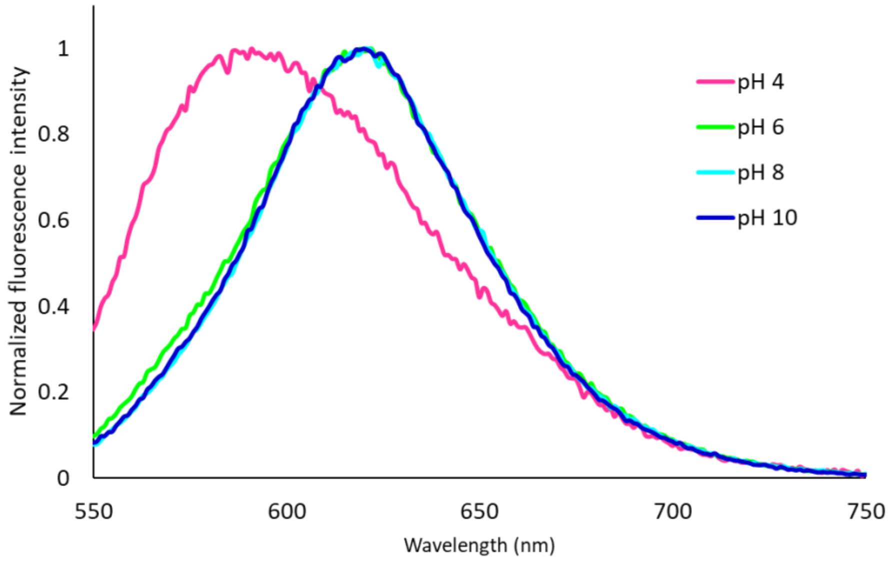


Figure S17: Normalized emission spectra of nile red in DDP suspension over varying pH at 45°C, when excited at 530 nm. N = 4. At pH 4, the emission maximum of nile red was found to be near 590 nm, whereas at pH 6, 8 and 10, the emission maximum was near 620 nm.

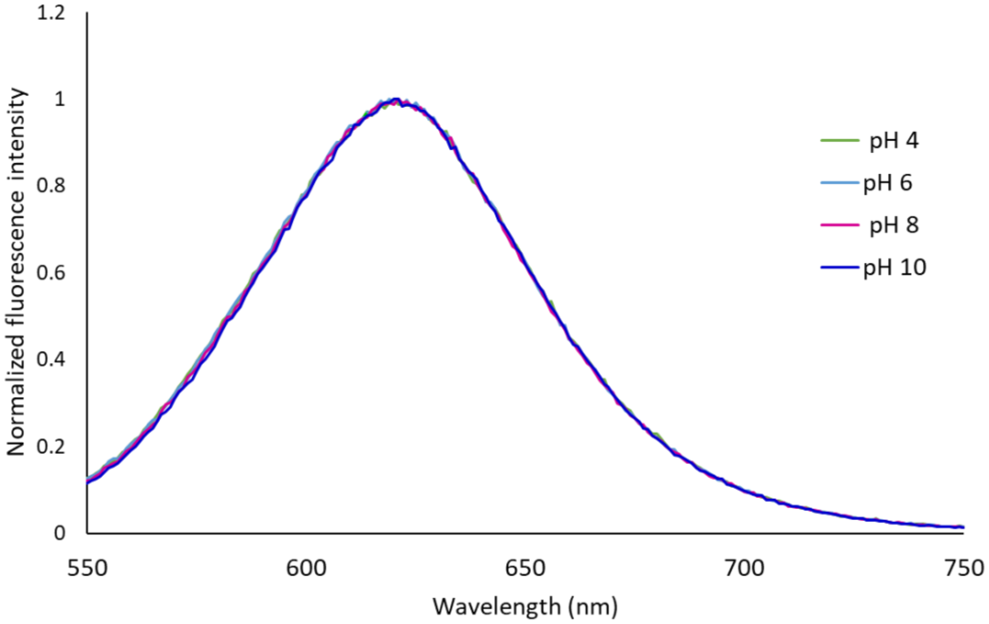


Figure S18: Normalized emission spectra of nile red in POPC (phospholipid) vesicle suspension over varying pH at 45°C when excited at 530 nm. N = 4. At all pH, the emission maximum of nile red was found to be near 626 nm. This showed that in POPC membranes, nile red emission maximum doesn't change with a change in pH.

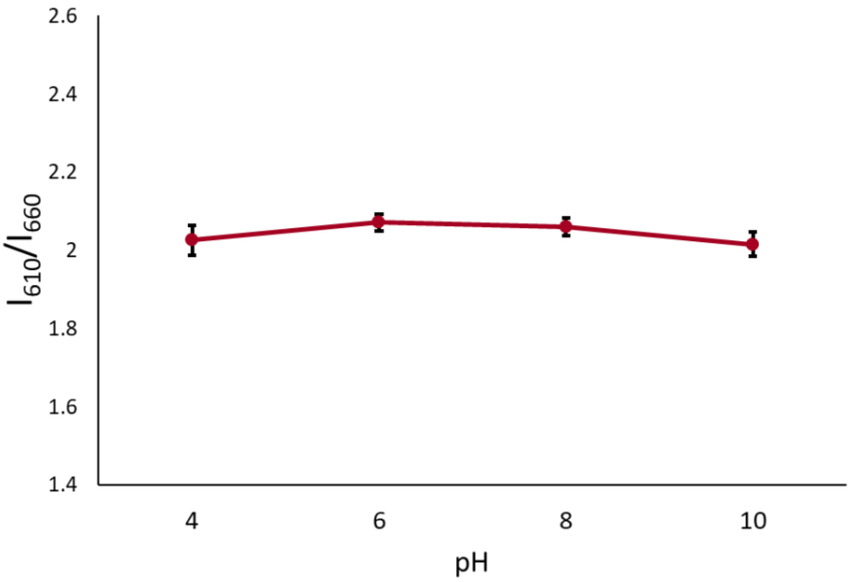


Figure S19: The scatter plot showing nile red $I_{610}/I_{660}$ intensity ratio of the POPC (phospholipid) vesicle suspension over varying pH at 45°C when excited at 530 nm. N = 4, error bar = SD. The $I_{610}/I_{660}$ remained unaltered (based on a two-tailed t-test) over varying pH.

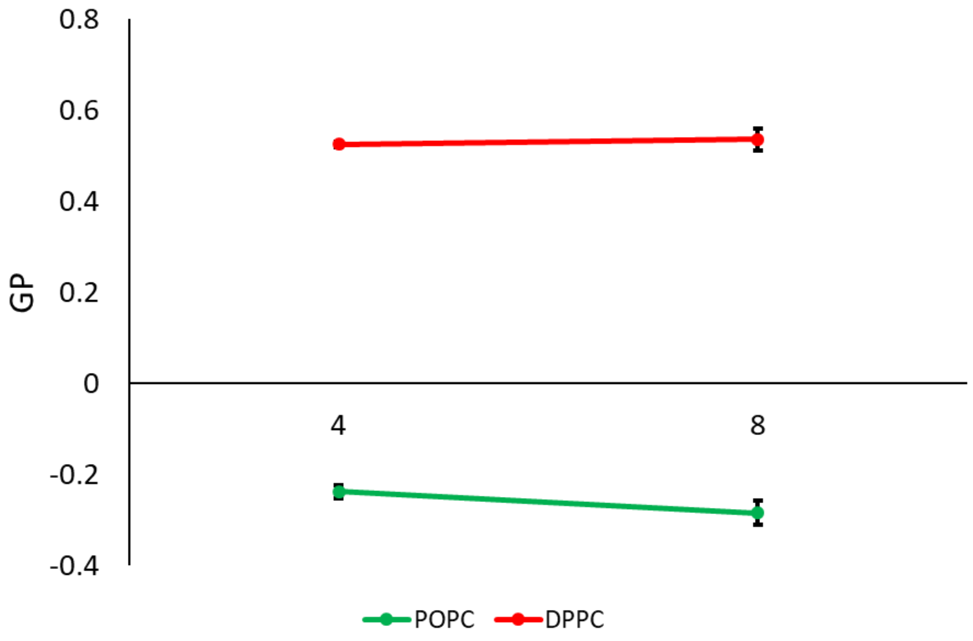


Figure S20: The scatter plot showing laurdan GP values of POPC and DPPC (phospholipids) vesicle suspensions over varying pH at 45°C when excited at 530 nm. N = 4, error bar = SD. The GP value remained unaltered (based on a two-tailed t-test) between the two pHs.

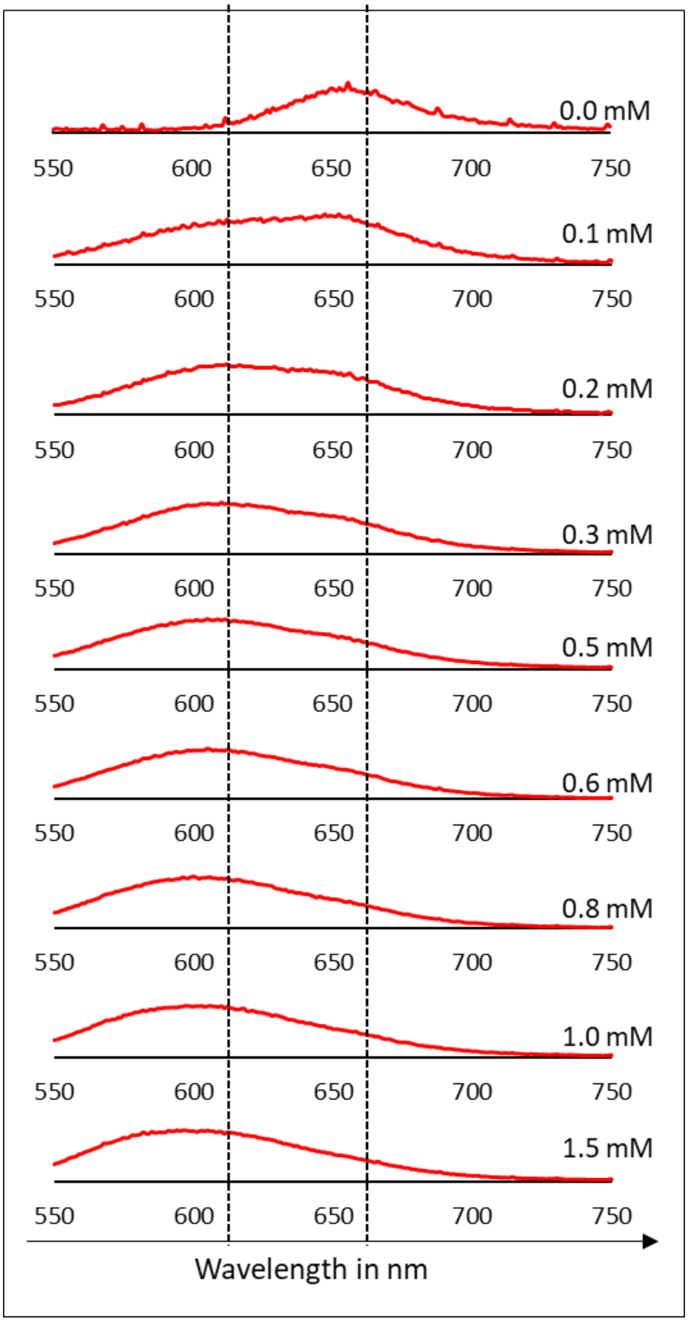


Figure S21: Normalized emission spectra of nile red in DDP suspension at pH 4 when plotted as the function of DDP concentration at 45°C. The black dotted lines indicate the emission at 610 and 660 nm; there is a blue-shift that occurred as a function of DDP concentration indicating an increase in hydrophobicity with increasing DDP concentration. N = 3.

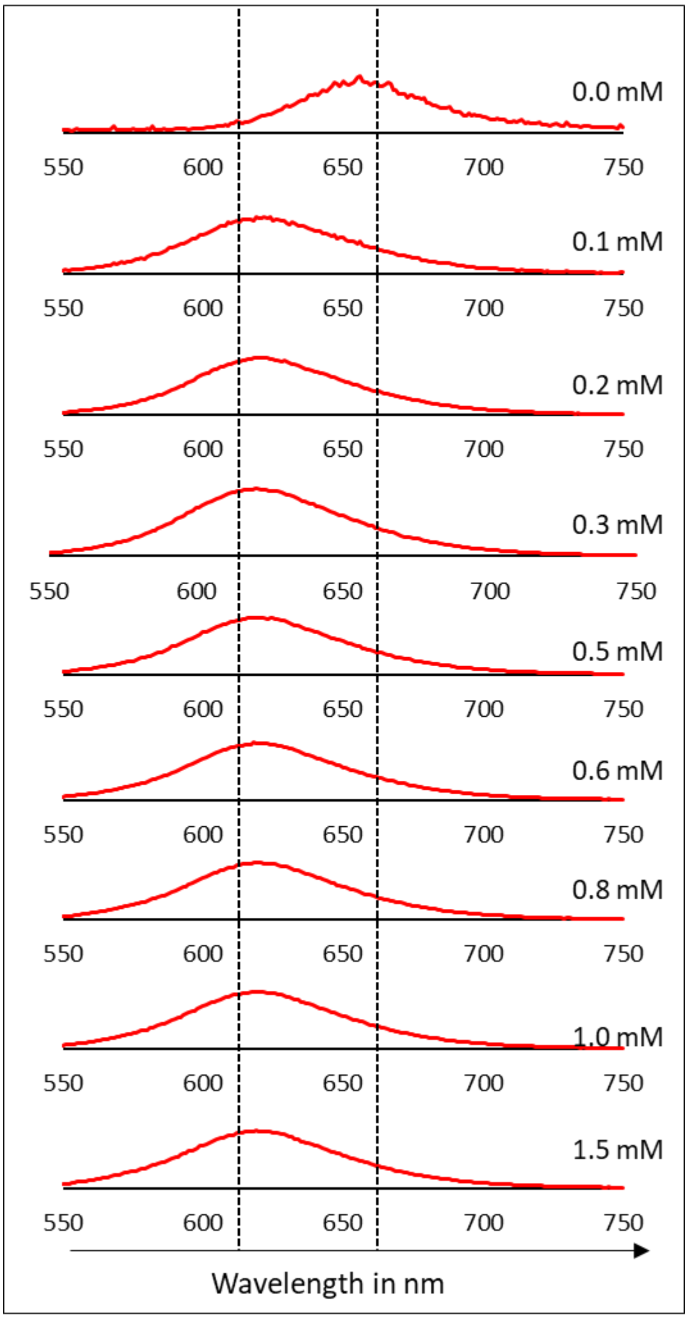


Figure S22: Normalized emission spectra of nile red in DDP suspension at pH 8 when plotted as the function of DDP concentration at 45°C. The black dotted lines indicate the emission at 610 and 660 nm, there is a drastic blue-shift upon addition of DDP (0.1 mM) with a maximum at 620 nm which remained same with further increasing DDP concentration. N = 3.

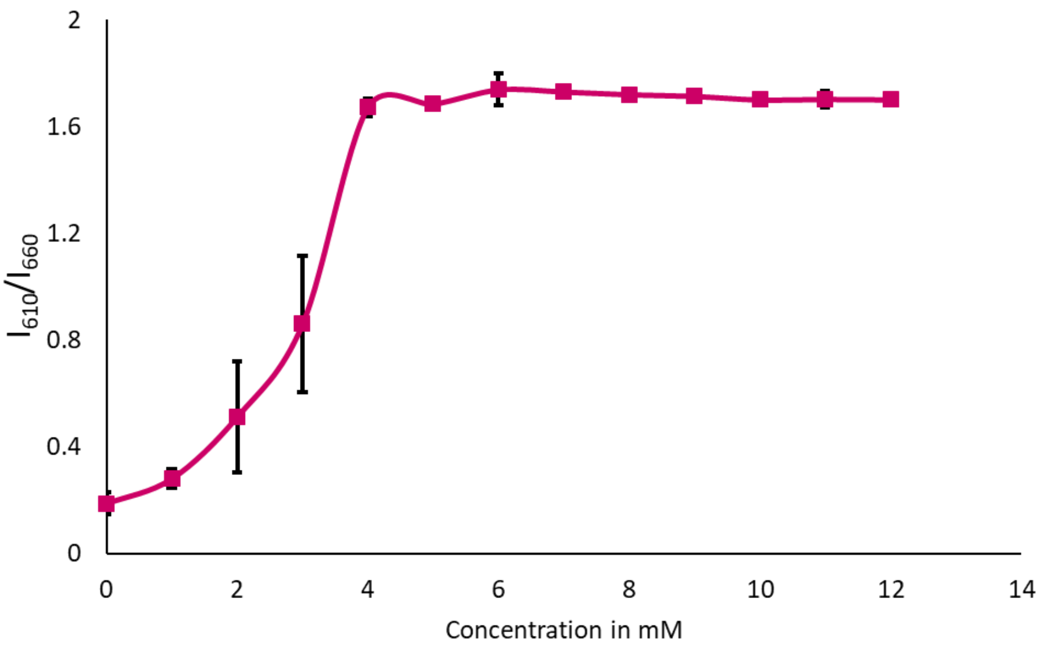


Figure S23: The scatter plot showing nile red $I_{610}/I_{660}$ intensity ratio of LA suspension at pH 8 when plotted as a function of DDP concentration. N = 3, error bar = SD. The $I_{610}/I_{660}$ was observed to saturate at around 6 mM indicating its CBC.

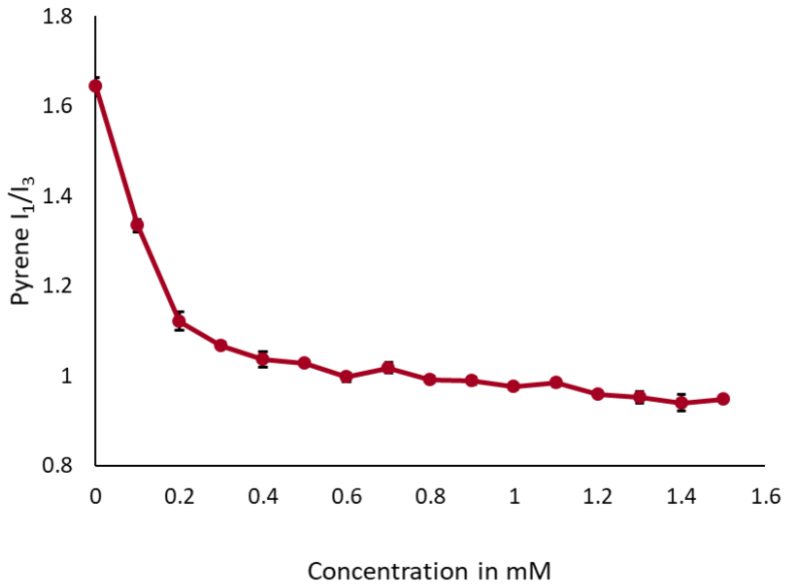


Figure S24: The scatter plot showing the pyrene $I_1/I_3$ ratio of DDP suspension at pH 4, plotted as a function of DDP concentration. N = 3, error bar = SD. The decay in $I_1/I_3$ was observed to saturate at 1±0.1 mM, indicating its CBC.

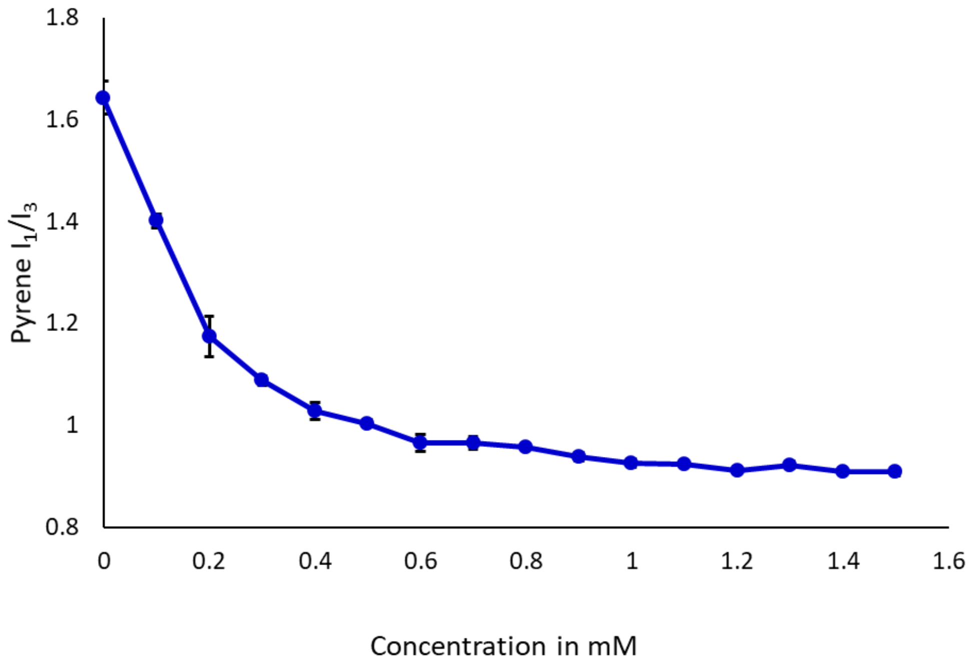


Figure S25: The scatter plot showing the pyrene $I_1/I_3$ ratio of DDP suspension at pH 8, plotted as a function of DDP concentration. N = 3, error bar = SD. The decay in $I_1/I_3$ was observed to saturate at 0.9±0.1 mM, indicating its CBC.

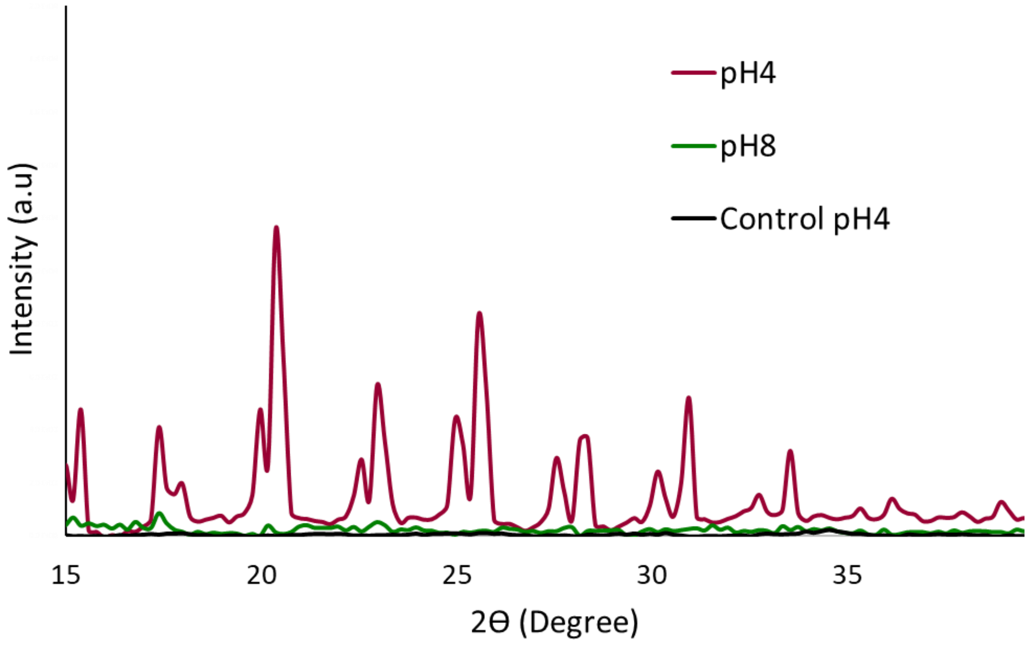


Figure S26: PXRD pattern of the dried film generated by repeatedly drop-casting and dried aqueous vesicle suspension of 10 mM DDP at pH 4 (magenta) and pH 8 (green), respectively. The black trace (control at pH 4) shows the XRD pattern of the dried film of extracted DDP molecules from suspension at pH 4 in chloroform:methanol::2:1. Sharp peaks were observed in the XRD pattern only at pH 4, depicting its crystalline nature. N = 4.

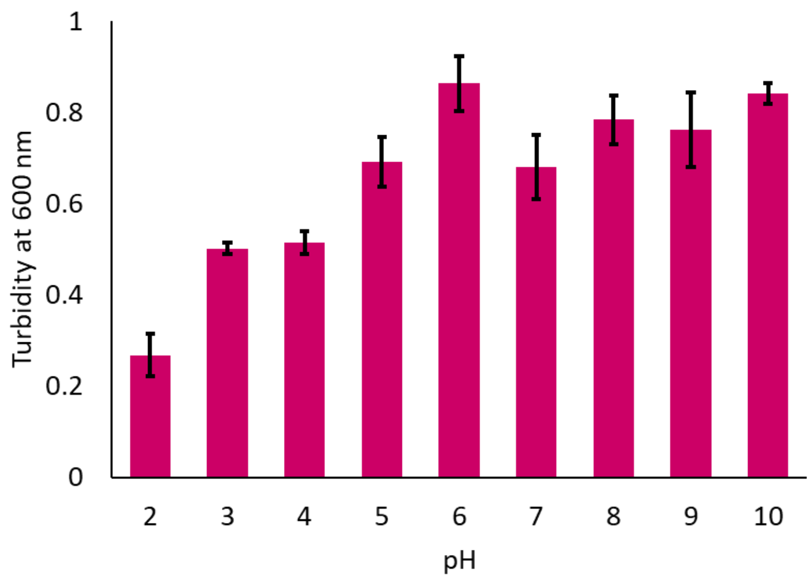


Figure S27: The bar plot shows the turbidity measurements at 600 nm for 10 mM DDP:DOH::1:1 membrane suspension at different pH (x-axis) and 45°C, N = 3, error bar = SD. Lowest turbidity was observed at pH 2 indicating comparatively less amount of higher order structures (vesicles and crystalline aggregates).

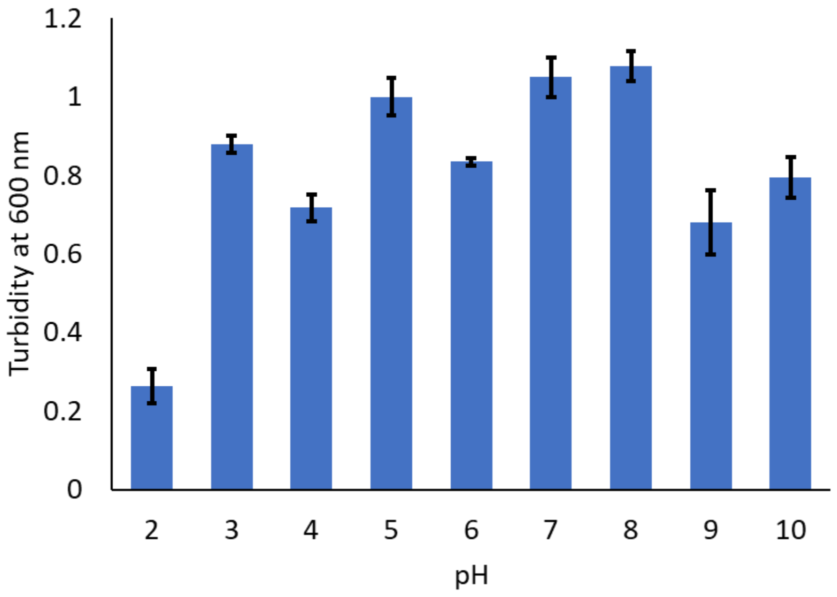


Figure S28: The bar plot shows the turbidity measurements at 600 nm for 10 mM DDP:DOH::2:1 membrane suspension at different pH (x-axis) and 45°C, N = 3, error bar = SD. Lowest turbidity was observed at pH 2 indicating comparatively less amount of higher order structures (vesicles and crystalline aggregates).

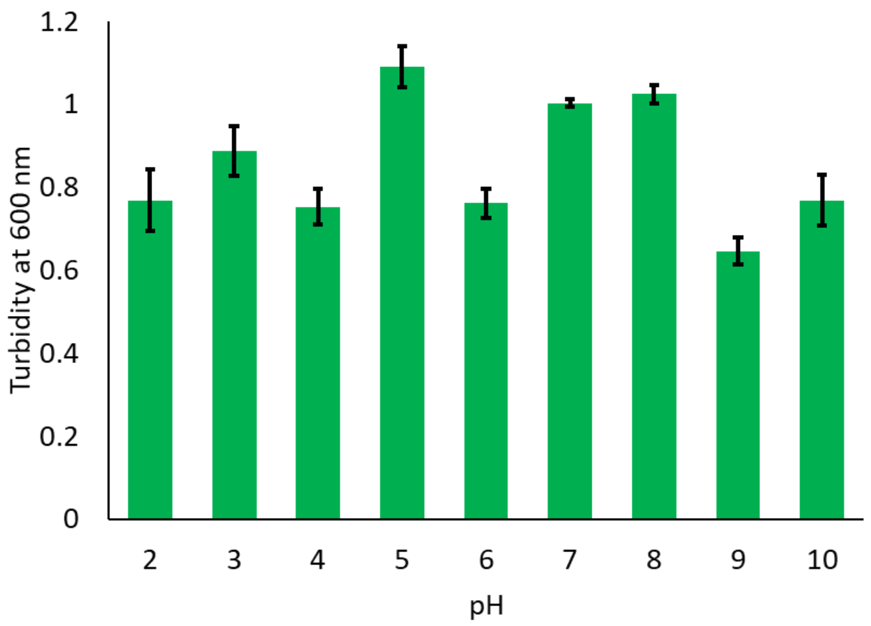


Figure S29: The bar plot shows the turbidity measurements at 600 nm for 10 mM DDP:DOH::4:1 membrane suspension at different pH (x-axis) and 45°C, N = 3, error bar = SD. Lowest turbidity was observed at pH 9 indicating comparatively less amount of higher order structures (vesicles and crystalline aggregates).

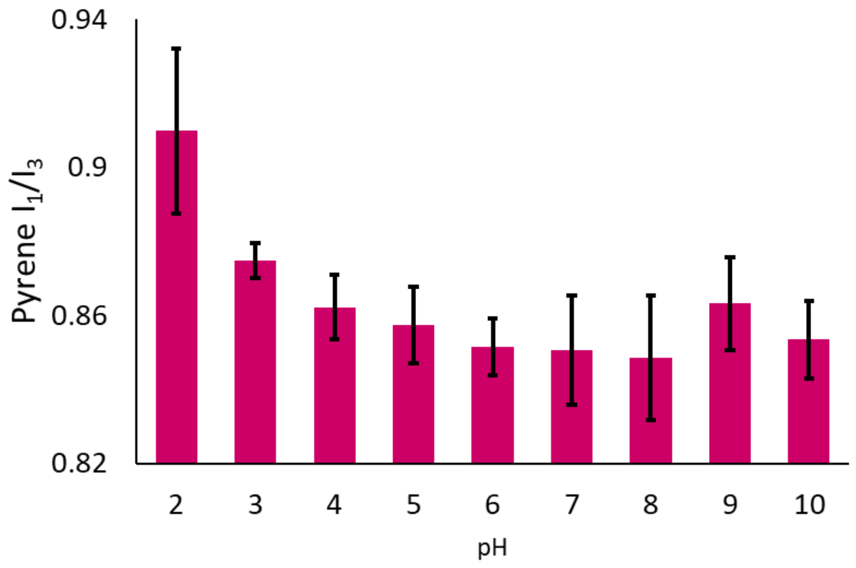


Figure S30: Pyrene $I_1/I_3$ ratio of 10 mM DDP:DOH::1:1 membrane suspension at different pH (x-axis) at 45°C, N = 4, error bar = SD. Higher ratio indicates less hydrophobicity (more water accessibility). The ratio was observed to be highest at pH 2. From pH 4-10, the ratio remains unchanged based on a two-tailed t-test.

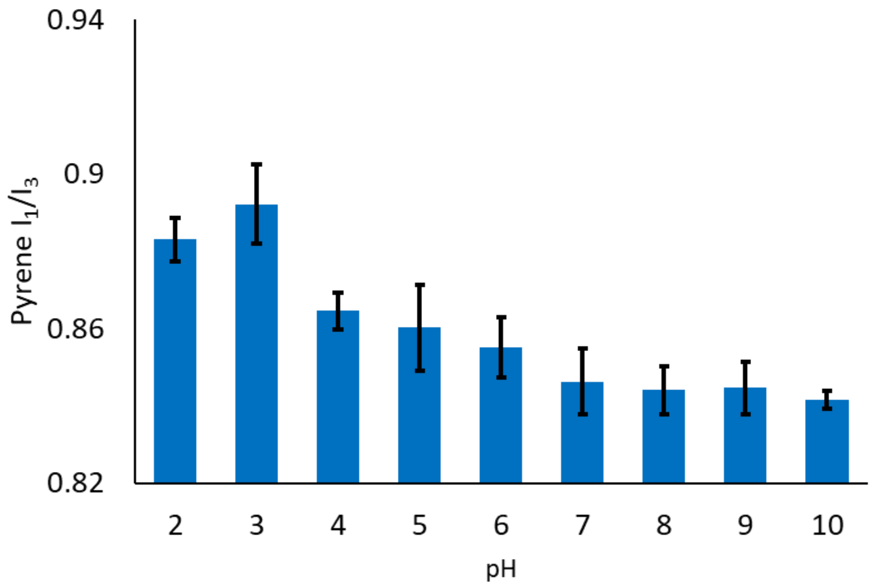


Figure S31: Pyrene $I_1/I_3$ ratio of 10 mM DDP:DOH::2:1 membrane suspension at different pH (x-axis) at 45°C, N = 4, error bar = SD. Higher ratio indicates less hydrophobicity (more water accessibility). The ratio was observed to be highest at pH 2-3. From pH 6-10, the ratio remains unchanged based on a two-tailed t-test.

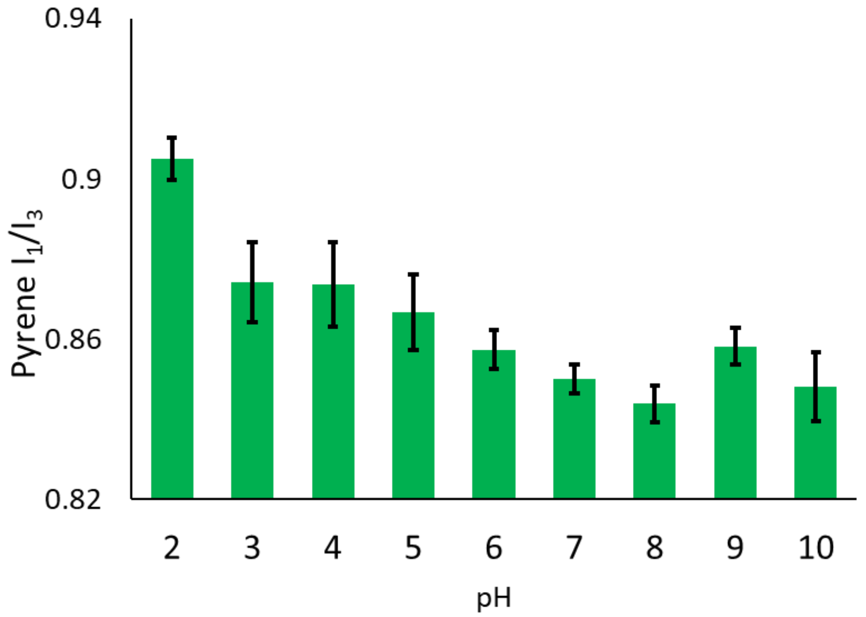


Figure S32: Pyrene $I_1/I_3$ ratio of 10 mM DDP:DOH::4:1 membrane suspension at different pH (x-axis) at 45°C, N = 4, error bar = SD. Higher ratio indicates less hydrophobicity (more water accessibility). The ratio was observed to be highest at pH 2 and lowest at pH 7-8.

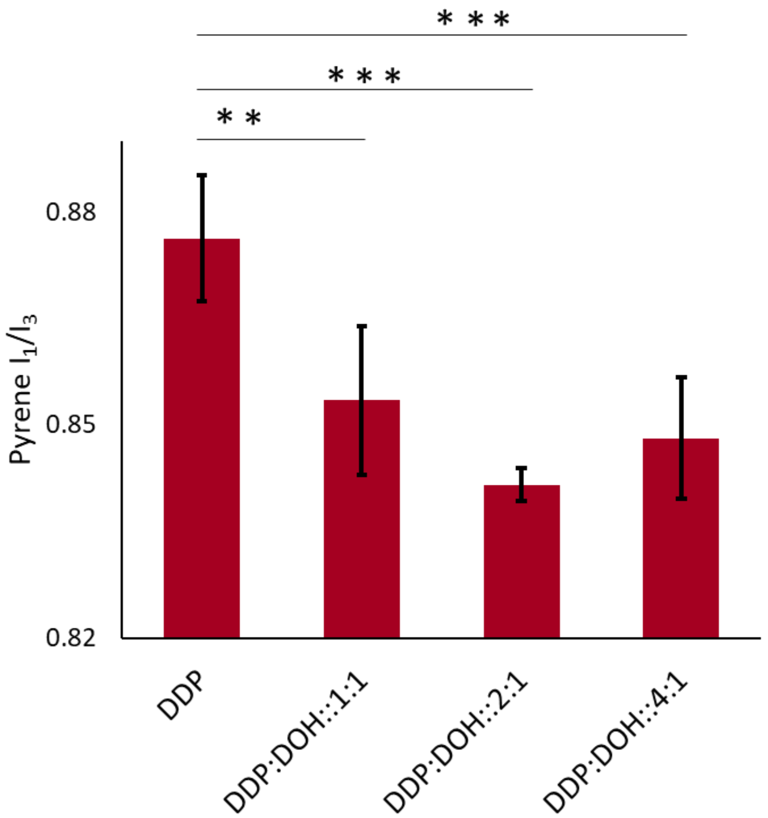


Figure S33: Pyrene $I_1/I_3$ ratio of 10 mM DDP and different DDP:DOH mixed systems at pH 10 recorded at 45°C, N = 3, error bar = SD. Upon comparison of only DDP membrane with the three DDP:DOH mixed systems, the decrease was found to be significant with a p-value of <0.05 based on a two-tailed t-test.

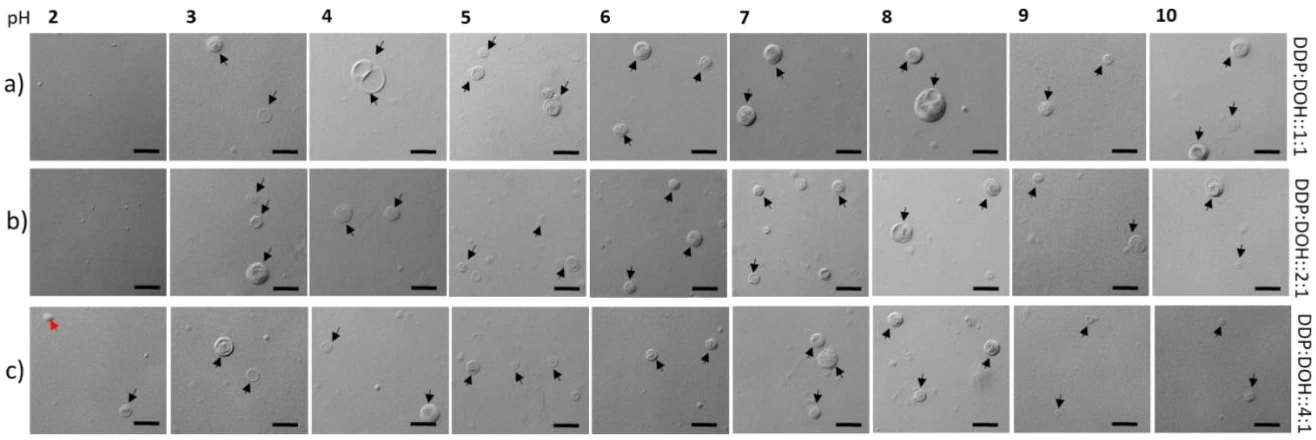


Figure S34: DIC microscopy images of three different DDP:DOH mixed membrane systems over varying pH. Panels (a) to (c) show three different systems, i.e., a) DDP:DOH::1:1, b) DDP:DOH::2:1 and c) DDP:DOH::4:1. Black and red arrows indicate vesicles and crystalline aggregates, respectively. N = 3, Scale bar = 10 µm. Vesicles were seen readily in most of the samples imaged across the varying pH excepting at pH 2 in DDP:DOH::1:1 and DDP:DOH::2:1 systems.

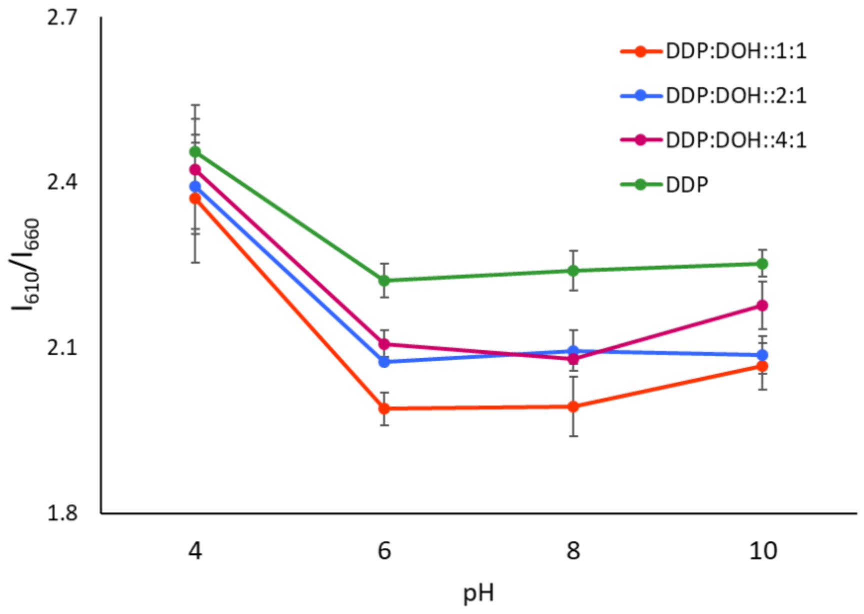


Figure S35: The scatter plot shows nile red $I_{610}/I_{660}$ ratio of the DDP membrane and the different DDP:DOH mixed systems over varying pH at 45°C. N = 4, error bar = SD. Higher $I_{610}/I_{660}$ ratio indicates low micropolarity. With an increase in pH, the ratio decreased showing an increase in micropolarity, for all the systems investigated. The lowest $I_{610}/I_{660}$ ratio was observed for DDP:DOH::1:1 system.

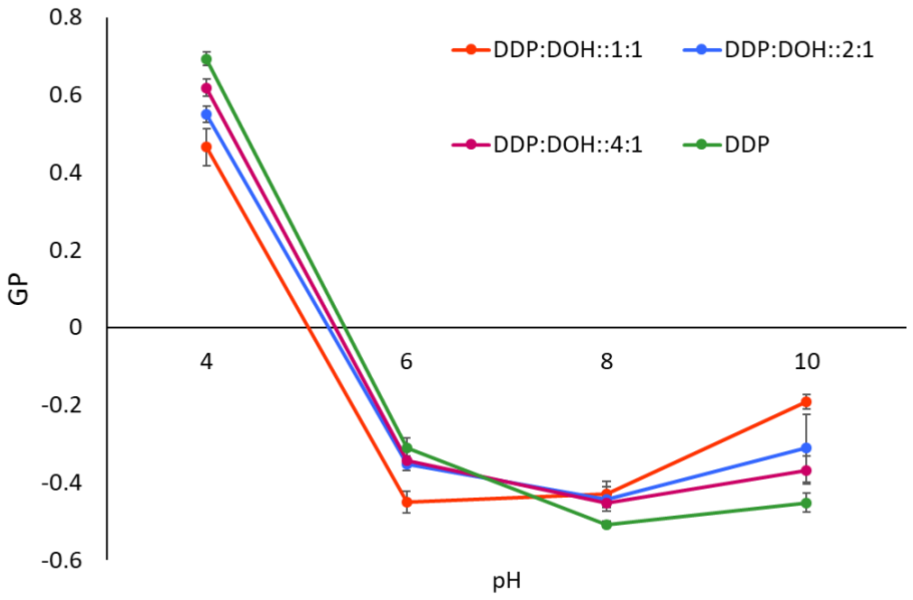

Figure S36:  The scatter plot shows laurdan GP values of the DDP membrane and the different DDP:DOH mixed systems over varying pH at 45°C. N = 4, error bar = SD. Higher GP value indicates higher membrane order. With an increase in pH, the GP value decreased showing a decrease in membrane order, for all the systems investigated.

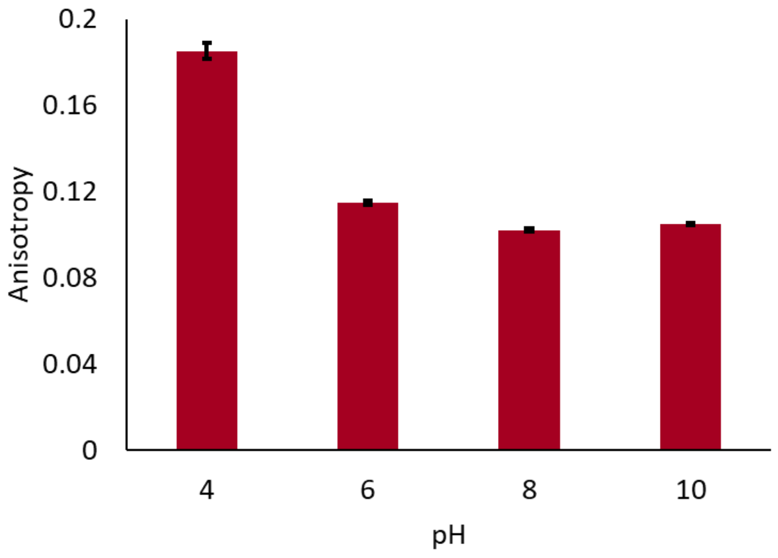


Figure S37: Laurdan anisotropy values of the DDP:DOH::1:1 membrane system over varying pH at 45°C. N = 4, error bar = SD. Higher Anisotropy value indicates lower fluidity. With an increase in pH, the anisotropy value decreased showing an increase in membrane fluidity.

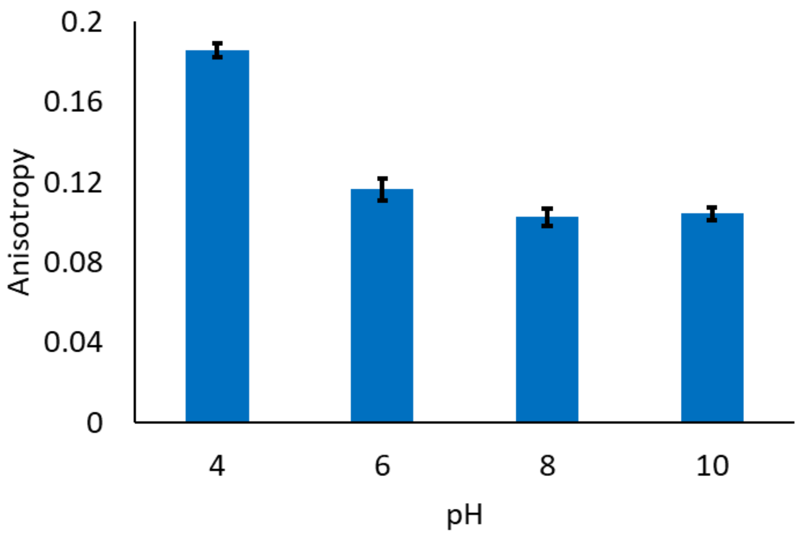


Figure S38: Laurdan anisotropy values of the DDP:DOH::2:1 membrane system over varying pH at 45°C. N = 4, error bar = SD. Higher Anisotropy value indicates lower fluidity. With an increase in pH, the anisotropy value decreased showing an increase in membrane fluidity.

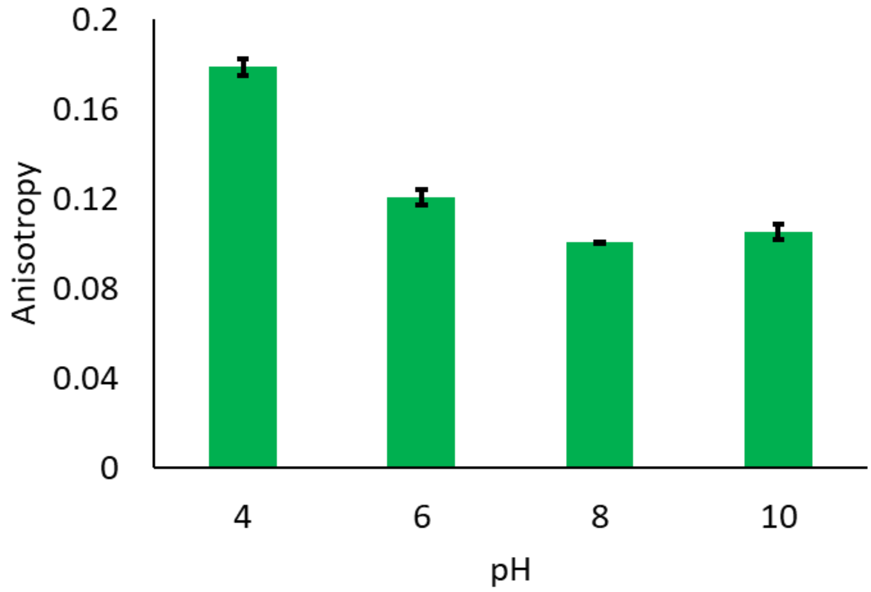


Figure S39: Laurdan anisotropy values of the DDP:DOH::4:1 membrane system over varying pH at 45°C. N = 4, error bar = SD. Higher Anisotropy value indicates lower fluidity. With an increase in pH, the anisotropy value decreased showing an increase in membrane fluidity.

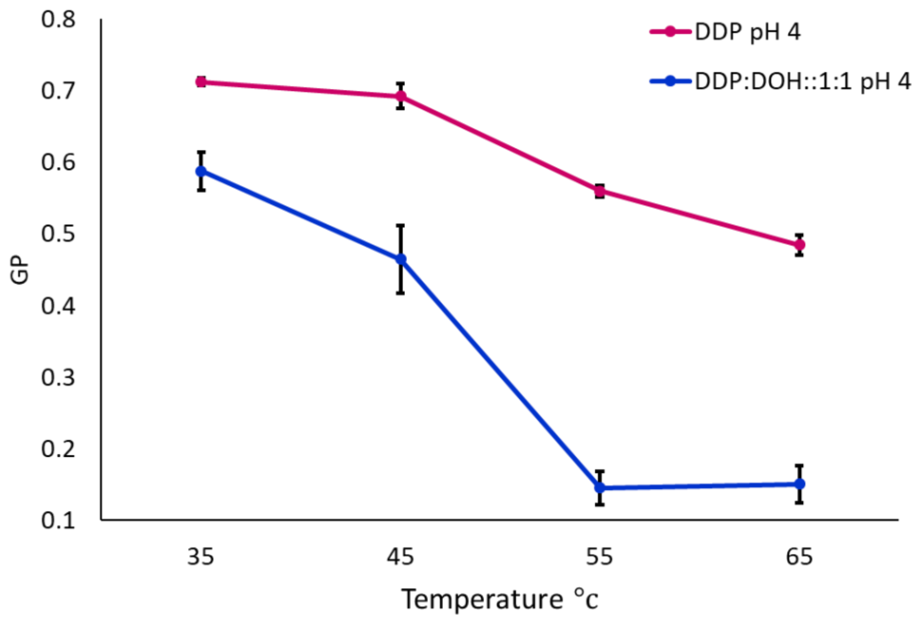


Figure S40: Graph showing laurdan GP values of the pure DDP and DDP:DOH::1:1 membrane systems over varying temperatures at pH 4. N = 4, error bar = SD. Higher GP value indicates higher membrane order. With an increase in temperature, the GP value decreased showing a decrease in membrane order, for both the systems. The extent of decrease was comparatively higher for DDP:DOH::1:1 system.

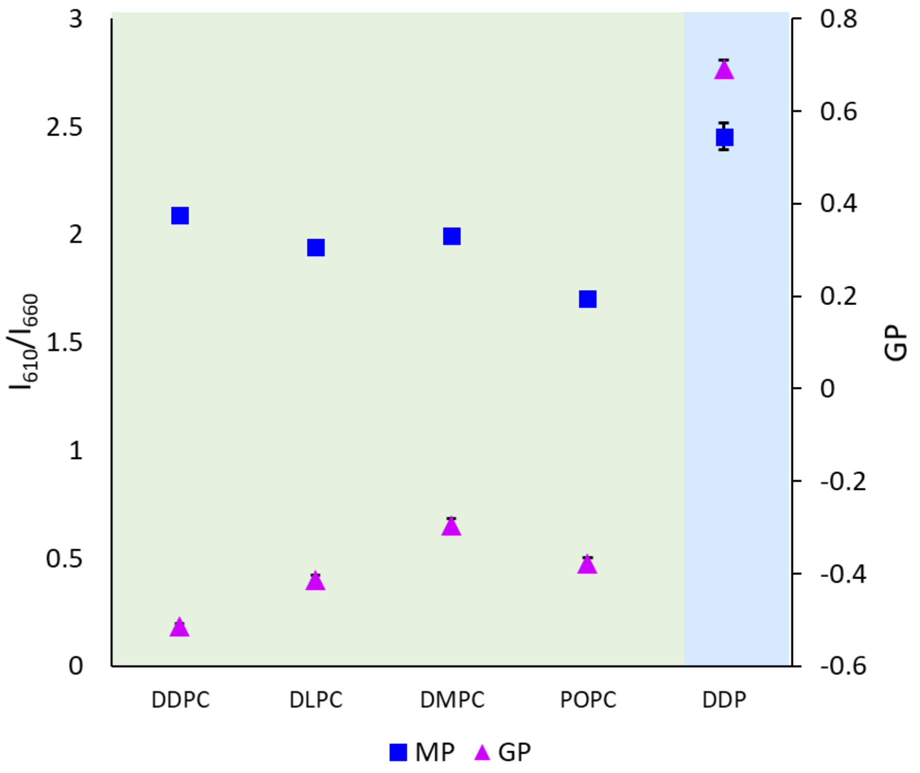


Figure S41: The scatter plot showing the nile red $I_{610}/I_{660}$ ratios (left y-axis) and the laurdan GP values (right y-axis) of different membranes i.e., for different phospholipids and DDP as indicated on x-axis, at pH 4 and 45°C. N = 4, error bar = SD. The $I_{610}/I_{660}$ ratio (MP) of DDP membrane was higher than all the four phospholipid membranes indicating lower membrane micropolarity. The GP value of DDP was found to be way higher than all the phospholipid membranes investigated indicating the tighter packing of DDP membrane.

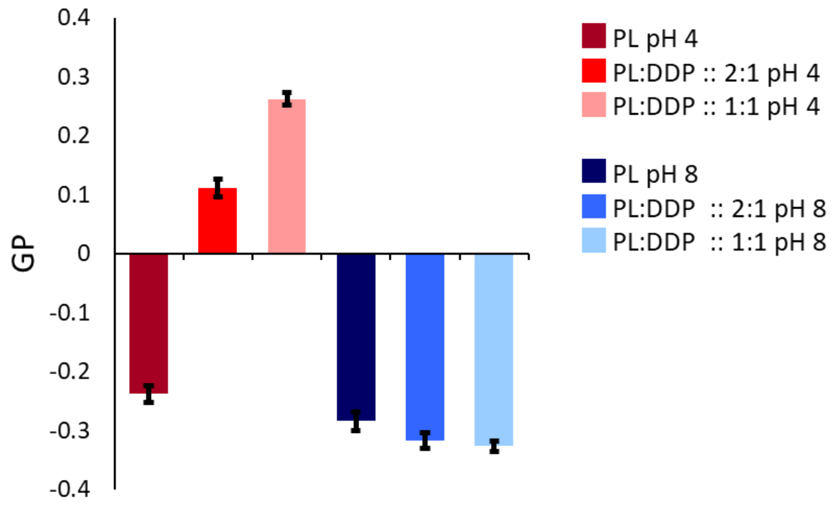


Figure S42: Bar graph showing laurdan GP values of DMPC (phospholipid) membranes and different DMPC:DDP mixed membranes at pH 4 and 8, respectively, at 37°C. N = 4, error bar = SD. Higher the GP value, higher the membrane order. At pH 4, the addition of DDP to PL membrane significantly increased its GP value indicating an increase in membrane order. Contrarily, at pH 8, the addition of DDP to PL membrane significantly decreased its GP value (based on a two-tailed t-test) indicating a decrease in membrane order.

Table ST1: The percentage of different protonated species (double protonated, mono protonated and deprotonated) of DDP (pka1 at 2.85 and pKa2 at 7.35) at different pH calculated using the Henderson-Hasselbalch equation.

| | Species percentage | | |
|---|---|---|---|
| **pH** | Double protonated **$C_{12}H_{25}OPO(OH)_2$** | Mono protonated **$C_{12}H_{25}OP(OH)O_2^-$** | Deprotonated **$C_{12}H_{25}OPO_3^{2-}$** |
| 2 | 87.5 | 12.49 | 6.25E-05 |
| 4 | 6.5 | 93.45 | 0.04 |
| 6 | 0.07 | 94.93 | 4.99 |
| 8 | 0.00 | 80.99 | 18.99 |
| 10 | 0.00 | 0.199 | 99.79 |

Table ST2: Laurdan GP values of different phospholipids (DMPC, POPC and DPPC) and the mixed systems composed of phospholipids and DDP in two different ratios (2:1 and 1:1) at pH 4 and 8, respectively, at 37°C. N = 4, SD = standard deviation.

| | | **GP (average)** | **S.D** |
|---|---|---|---|
| **DMPC** | PL pH 4 | -0.14 | 0.01 |
| | PL:DDP :: 2:1 pH 4 | 0.24 | 0.02 |
| | PL:DDP :: 1:1 pH 4 | 0.43 | 0.02 |
| | PL pH 8 | -0.16 | 0.02 |
| | PL:DDP :: 2:1 pH 8 | -0.19 | 0.01 |
| | PL:DDP :: 1:1 pH 8 | -0.21 | 0.01 |
| **POPC** | PL pH 4 | -0.23 | 0.01 |
| | PL:DDP :: 2:1 pH 4 | 0.11 | 0.01 |
| | PL:DDP :: 1:1 pH 4 | 0.26 | 0.01 |
| | PL pH 8 | -0.28 | 0.01 |
| | PL:DDP :: 2:1 pH 8 | -0.31 | 0.01 |
| | PL:DDP :: 1:1 pH 8 | -0.32 | 0.00 |
| **DPPC** | PL pH 4 | 0.52 | 0.00 |
| | PL:DDP :: 2:1 pH 4 | 0.64 | 0.00 |
| | PL:DDP :: 1:1 pH 4 | 0.65 | 0.01 |
| | PL pH 8 | 0.53 | 0.02 |
| | PL:DDP :: 2:1 pH 8 | -0.00 | 0.02 |
| | PL:DDP :: 1:1 pH 8 | -0.11 | 0.02 |
